# Continuous, multidimensional coding of 3D complex tactile stimuli by primary sensory neurons of the vibrissal system

**DOI:** 10.1101/869255

**Authors:** Nicholas E. Bush, Sara A. Solla, Mitra J. Z. Hartmann

## Abstract

To reveal the full representational capabilities of sensory neurons, it is essential to observe their responses to complex stimuli. In the rodent vibrissal system, mechanical information at the whisker base drives responses of primary sensory neurons in the trigeminal ganglion (Vg). Studies of how Vg neurons encode stimulus properties are typically limited to 2D analyses and restricted stimulus sets. Here we record from Vg neurons during 3D stimulation while quantifying the complete 3D whisker shape and mechanics. Results show that individual Vg neurons simultaneously represent multiple mechanical features of the stimulus, do not preferentially encode principal components of the stimuli, and represent continuous and tiled variations of all available mechanical information. As a population, the neurons span a continuum of rapid and slow adaptation properties; a binary distinction between these adaptation classes is oversimplified. These results contrast with proposed codes in which Vg neurons segregate into functional classes.

## Introduction

Sensory neuroscience aims to quantify how neurons encode and process fundamental physical stimuli: photons, pressure waves, chemicals, and mechanical forces. A common experimental approach is to use controlled, reduced, and repeatable stimulus sets to elicit consistent neural responses that can be averaged to reduce trial-to-trial variability^1-6^.This method lends itself to a description of neural coding in which neurons are tuned to a small number of stimulus features^7^, and to a categorization of neurons into functional classes^8^ based on their differential responses to the stimuli^1,3,5,9-12^. A problem with this approach is that results are constrained by the stimuli; these are typically chosen to be categorical and significantly underrepresent the stimulus space to which the neurons respond. It is thus almost inevitable that the neurons will, in turn, exhibit simple, low-dimensional tuning curves and categorical response types. Descriptions of neural representations of stimuli therefore remain incomplete.

The rodent whisker system is one of the premier models for studying tactile processing and cortical function^13^. During natural exploration, rodents move their whiskers in rhythmic, non-repeatable 3D trajectories^14,15^ generating complex, continuously varying patterns of tactile input. However, most descriptions of vibrissal-responsive primary sensory neurons in the trigeminal ganglion (Vg) are based on experiments that use reduced stimulus sets, in which variations in the presented stimuli are discrete, involving only a few features or limited spatial directions, small in dynamic range, or presented only along the neuron’s preferred direction. Recent work in awake, whisking animals has employed a more continuous stimulus set, but these studies have been restricted to a 2D analysis of whisker motion^16-19^.

Here we take inspiration from studies of the visual system that employ increasingly complex stimulus sets^20,21^, and apply to the whiskers a manual, naturalistically varying stimulus set designed for a rich exploration of tactile stimulus space. The stimuli employed here span ranges similar to those observed in naturally behaving animals^22,23^, Supplementary Fig. 2). We introduce a stereo-vision 3D whisker imaging technique, apply a model of 3D whisker mechanics^24^, and implement recently developed statistical modeling techniques^25,26^ to characterize the full input space available to the whisker system and the consequent response properties of Vg neurons.

When characterized through the expanded, naturalistic stimulus set employed here, the response properties of Vg neurons reveal a fundamentally different encoding structure than generally appreciated. We find that Vg neurons are broadly tuned across multiple stimulus features, including force, bending moment, and rotation, as well as stimulation direction. These neurons do not represent select mechanical features, nor do they represent the structure of the leading principal components that span a low dimensional subspace of the mechanical stimuli space. Importantly, these diffuse representations of mechanical stimuli continuously tile the relevant stimulus space, suggesting that primary sensory neurons possibly use a dense coding scheme to represent tactile stimuli. Thus, Vg neurons do not segregate into functional classes that convey specialized feature information to more central structures^4,27,28^.

An important lesson conveyed by these results is the crucial fact that descriptions of the response properties of sensory neurons are fundamentally constrained by the complexity and extent of the stimulus sets used to probe them, and that stimulus sets that underrepresent the extent and complexity of natural stimuli can lead to incomplete, if not incorrect, descriptions of the encoding properties of these neurons.

## Results

### Acquisition of 3D stimulus information

We recorded 78 whisker-responsive Vg neurons in 22 anesthetized rats during manual tactile stimulation of single whiskers. During stimulation, high-speed video (300 or 500 fps) captured whisker motion in two views (Fig. 1A). Following previous methods^29^, a graphite probe was used to repeatedly deflect the whisker at 2-3 different distances along its length (Fig. 1C) in 8 cardinal directions (Fig. 1D). Stimulation speed varied across trials, with two distinct speeds (“fast” and “slow”) at each stimulation location. Video and neural data were recorded for an average of ∼500s per neuron, with an average of 684 whisker deflections across all conditions. Whiskers were tracked in both camera views^30^, and the 3D whisker shape and stimulus contact point reconstructed (Fig. 1B). Mechanical models^24^ were used to compute forces and moments at the whisker base for each video frame (Fig. 1E).

**Figure 1:**
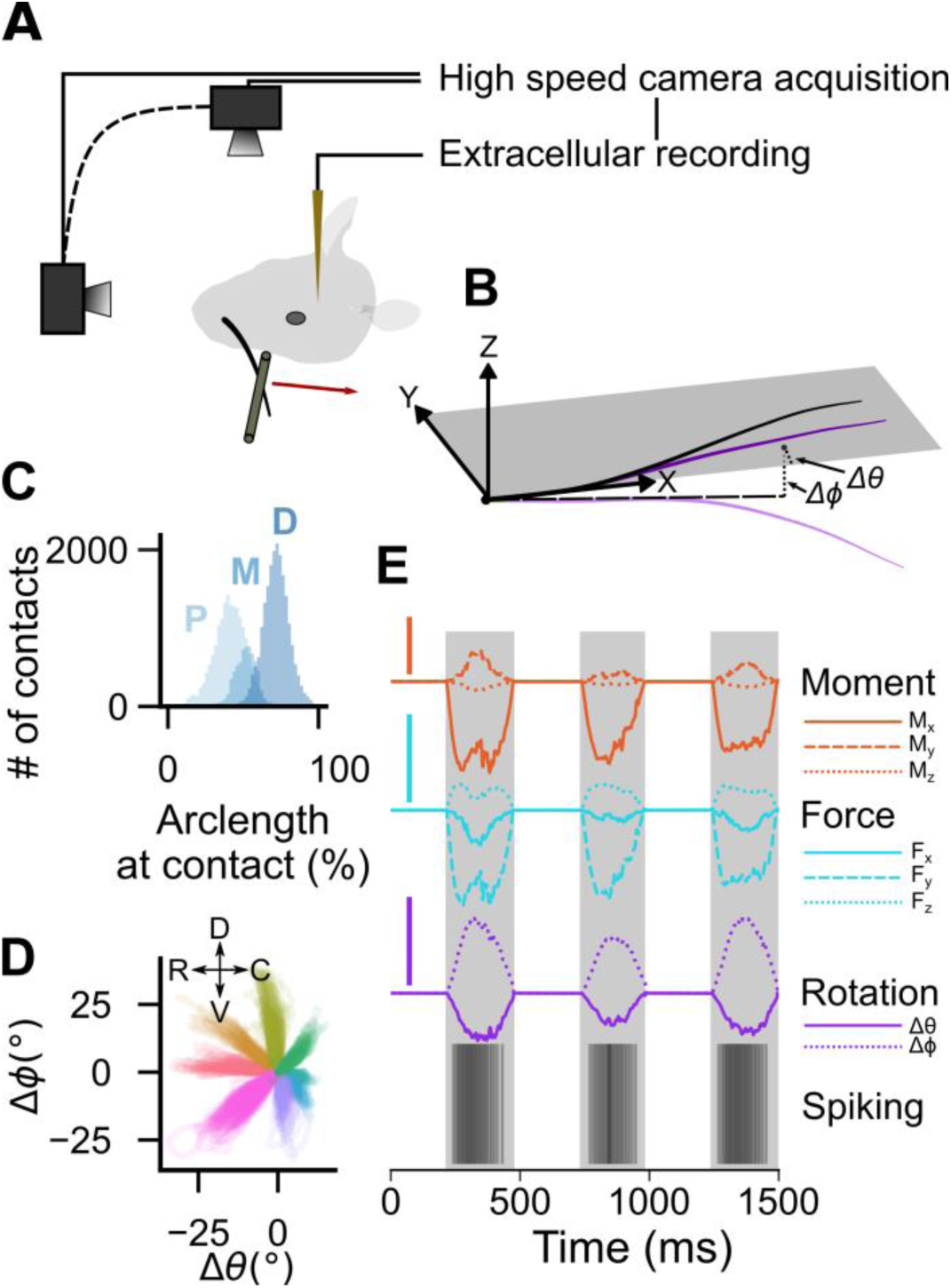
Acquisition of 3D stimulus information. **(A)** Schematic of experimental setup. A tungsten electrode records activity from a Vg neuron as manual deflections of a single whisker are monitored with two high-speed cameras. **(B)** Example 3D reconstruction of a whisker. Mechanics are calculated in whisker-centered coordinates based on whisker shape changes (bending; yellow). Rotational features are calculated based on changes in base segment emergence angles (green, Δ*θ* in the x-y plane, Δ*ϕ* in the x-z plane) compared to rest (black). **(C)** Deflections were applied at 2 or 3 distances along the whisker arclength; whisker arclengths were normalized to one and contacts sorted into three “distance groups”: proximal (P), middle (M), and distal (D). A histogram of the number of contacts at a given arclength is shown across all neurons. When only two distinct clusters were found for a given whisker, the middle group was omitted and is thus underrepresented. **(D)** Deflections were applied in approximately 8 directions in the plane perpendicular to the whisker’s primary axis and deflections were sorted into eight “direction groups.” Trajectories described in terms of the two base angles are shown for all deflections of an example whisker; color indicates the assigned direction group. Qualitatively distinct groups are observed for this example and for all whiskers. **(E)** Traces of moments (orange; *M*_*x*_, *M*_*y*_, *M*_*z*_), forces (cyan; *F*_*x*_, *F*_*y*_, *F*_*z*_), rotation angles (magenta; Δ*θ*, Δ*ϕ*), and observed spikes for three successive deflections are shown for an example whisker/neuron pair. Scale bars are 0.1 *μNm*, 0.5 *μN*, and 5° respectively.

In the anesthetized animal, exerting a force on a whisker causes the whisker to bend and the follicle to rotate within the mystacial pad^29^. This rotation generates a force between the follicle and surrounding tissue that may contribute to Vg responses. Because the mechanical properties of the follicle-cheek interface are unknown, we used the whisker’s angular rotation as it emerged from the cheek (Fig. 1E, Supplementary Fig. 1) as a proxy for the force of the tissue on the follicle. All signals, including spike times, were interpolated and binned at 1 kHz.

The variability inherent in manual deflections precludes comparing time-locked responses across trials. However, each manual deflection evolved similarly over time, as would occur during repeated whisks against an object. This temporal structure lends itself to “time-normalized” analyses in which stimulation durations are normalized to one^16,28^.

### Most neurons are jointly tuned to direction and location of stimulation along the whisker arclength

Previous studies demonstrated that Vg firing rate is strongly influenced by both deflection direction^2,10,12,31-33^ and the arclength of stimulus contact^4,28^. However, these studies have not quantitatively examined the joint effect of these parameters. We quantified the effects of simultaneous changes in both arclength of contact (two or three groups) and direction (eight groups). For each neuron, we computed the average firing rate across many deflections for each arclength/direction combination, and used it to compute a Directional Selectivity Index (DSI) defined as (1 − *σ*^2^), where *σ*^2^ is the directional circular variance^34^ (see *Methods*).

Results of this analysis are shown in Figures 2A-D for one example neuron. This neuron’s firing rate increased as stimulation became increasingly proximal (Fig. 2A,C. 2-way ANOVA F=509 main effect of arclength, p<0.001; Tukey’s post-hoc test p<0.05). The neuron’s preferred direction was near 225° (Fig. 2B,C; F=305 main effect of direction, p<0.001;Tukey’s post-hoc test: p<0.05 for 22/28 multiple comparisons), and it exhibited a moderate DSI (0.54 for proximal stimulation, 0.64 for distal stimulation) (Fig. 2D). For each cell, the DSI was calculated by bootstrapping; we randomly sampled half of all contacts without replacement for 1000 replicates to compute the mean DSI and its standard deviation. The example neuron was more directionally tuned for distal contacts (Student’s t-test: t=123, p<0.001). Notably, multiple combinations of arclength and direction can result in the same firing rate (Fig. 2C).

**Figure 2:**
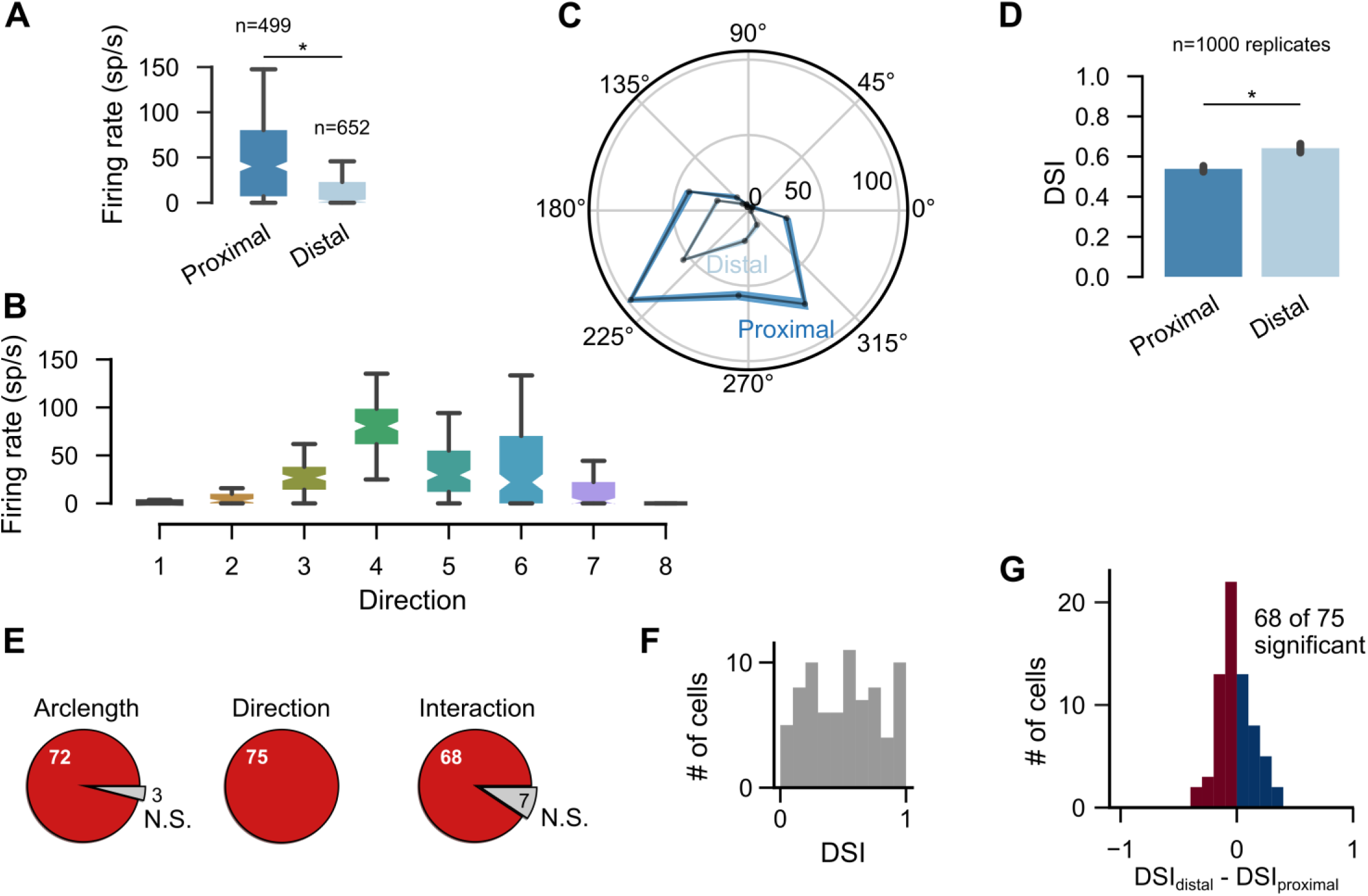
Most neurons have firing rates that correlate both with arclength of contact and deflection direction: **(A-D)** An example neuron whose firing rate is modulated by both arclength and direction. In A and B, boxes are median +/- 1 quartile and capped lines are median +/- 2 quartiles. **(A)** Firing rate increases as stimulation becomes more proximal (*p>0.001). **(B)** Firing rate modulation by direction group. The DSI for this neuron is 0.72. **(C)** Average firing rate and directional selectivity can depend on the arclength of contact. Average firing rate of the example cell is shown as a function of deflection direction, for both proximal and distal arclength of contact. The lines indicate average firing rate for all deflections in each group, with line thickness indicating +/- S.E.M. Note that for this neuron the directional tuning curve is broader for more proximal stimulation, indicating a weaker directional tuning for more proximal deflections. **(D)** Quantification of the DSI for the example cell. Bar is mean +/- S.D. (*p<0.001). **(E)** Number of cells with significant effects of arclength, direction, and their interaction on firing rate (two-way ANOVA). Red indicates significant effects. **(F)** Vg neurons range from not at all directionally modulated (DSI=0), to very strongly modulated (DSI=1). **(G)** Directional tuning strength is modulated by arclength of contact for most cells. Cells with positive (DSI_distal_ – DSI_proximal_) are more directionally tuned for distal contacts (28/68 neurons).

These results are generalized over all neurons in Figures 2E,F. Of the 78 recorded neurons, 75 had distinguishable arclength and direction groups. Although all 75 neurons exhibited significant direction tuning (two-way ANOVA p<0.05), the DSI was continuously and uniformly distributed across all neurons (Fig. 2F; Kolmogorov-Smirnov test: n=75, D=0.068, p=0.88), revealing a continuum of directional tuning strength across the population. Nearly all neurons (72/75) were also tuned for arclength, with proximal stimulation typically evoking stronger responses.

Importantly, the firing rate of most neurons (68/75, two-way ANOVA, p<0.05) was modulated by both direction and arclength. Figure 2G shows the change in DSI for distal compared to proximal stimulations for all cells. Cells with (DSI_distal_ – DSI_proximal_) greater than zero were more directionally tuned for distal contacts (28/68 neurons). Note that approximately equal numbers of neurons become more/less directionally tuned as stimulation became increasingly distal

The results of Figure 2 indicate that naturalistic, complex stimulation can recapitulate classical Vg responses observed during controlled ramp-and-hold stimulation^3,9,10^. Figure 2 also accentuates an underappreciated characteristic of Vg responses: when direction and arclength covary continuously and simultaneously, as during natural contact, Vg firing rate is governed jointly by both parameters. These results suggest that a single neuron’s response cannot unambiguously encode a stimulus feature, and that a population readout is required.

### Temporal patterns of spikes during contact are complex and direction dependent

The previous analyses characterized average Vg firing rates, but detailed temporal features of Vg spike patterns are important in shaping central responses^35^. We exploited variability across individual deflections to quantify the dependencies between temporal firing pattern and deflection direction across cells (Supplementary Videos 1-3). Two examples are shown in the time-normalized histograms of Figure 3A. Cell 1 exhibits strong changes in average firing rate with stimulation direction, with no change in temporal firing pattern. In contrast, the firing pattern of Cell 2 varies significantly with stimulation direction; some directions show a strong onset response, others a strong offset response, and yet others show neither.

**Figure 3:**
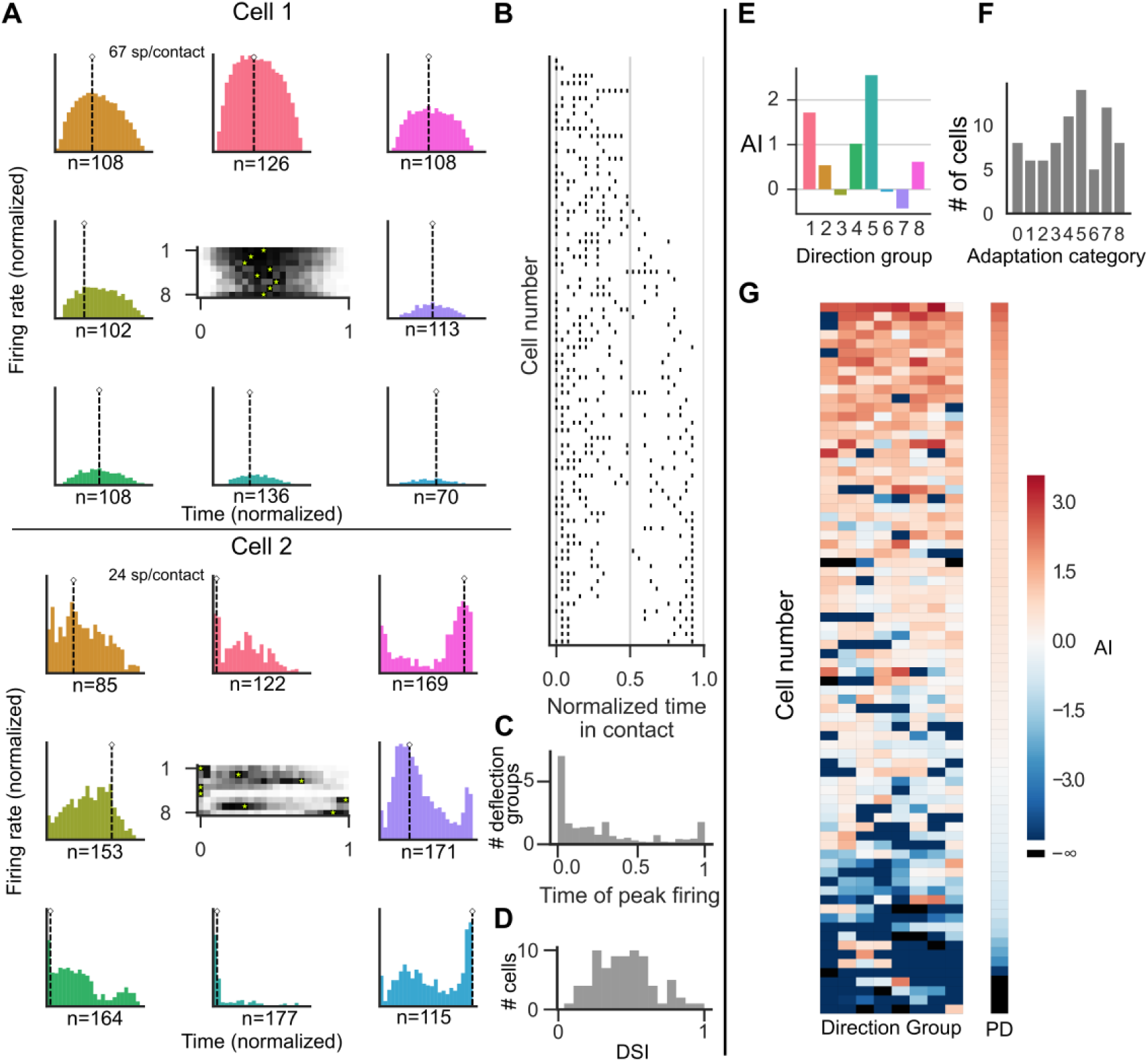
The temporal structure of Vg responses can be complex and direction dependent. **(A)** Normalized peristimulus time histograms (PSTHs) grouped by deflection direction for two example neurons. Horizontal axes are normalized to a contact duration between 0 and 1. Vertical axes are scaled up to the neuron’s maximum firing rate; direction groups are color-coded as in Figure 1D. Color, position, and number denote direction group of the PSTH. The number of contacts (n) used to construct each PSTH is indicated below each horizontal axis. Cell 1 shows little change in temporal pattern with deflection direction, while Cell 2 shows significant modulation. Central plots show the eight histograms; y-axis indicates direction group. Firing rate is shown in grayscale; a yellow asterisk indicates peak firing rate, which can occur between contact onset (time = 0) and offset (time = 1). **(B)** Time of peak firing rate(s) for all neurons in all directions. Each row represents a neuron; each row has eight points, one for each direction group. Identical peak times for several directions appear as superimposed points. Cells are ordered by variance of peak time of firing. **(C)** Histogram of time of peak firing collapsed across neurons and directions. **(D)** DSI of peak time as a function of direction across neurons. If peak time is heavily modulated by direction, DSI≅1. **(E)** Adaptation index (AI) for each direction for Cell 2. (**F**) Adaptation category for all cells. **(G)** AI for all neurons and direction groups. Cells are ordered by mean AI across directions. Right column (PD) isolates the AI for each neuron’s preferred direction, ordered by decreasing AI.

The center plots in the two examples of Figure 3A show time-varying firing rates as grayscale heatmaps for each direction, with the times of maximal firing rate (“peak times”) indicated by yellow asterisks. The peak times for all cells and directions are shown in Figure 3B. Each cell has eight peak times, one per direction. Cells are ordered by peak time variance; cells with little directional modulation of peak time are at the top and those with strong modulation at the bottom. Figure 3C aggregates data across neurons and directions; peak times are most likely to occur at onset, less likely at offset, and least likely in the middle of deflection. The influence of deflection direction on the peak time was quantified with the DSI of the peak time; Figure 3D shows that the DSI of peak time is normally distributed (Shapiro Test: n=77, W=0.98, p=0.49). Deflection direction has a moderate influence on spiking patterns, with few neurons very strongly (DSI ≅ 1) or very weakly (DSI ≅ 0) modulated.

Vg neurons are frequently classified as rapidly adapting (RA) or slowly adapting (SA) based on their response to ramp-and-hold stimulation^3,9-11,32^. To perform a similar analysis with the present data we introduce an “Adaptation Index” (AI) as the log ratio of the firing rate during the first 10 ms of contact to the average firing rate. The AI is calculated separately for each direction group. An AI of 0 indicates no difference between onset firing rate and mean firing rate; a strongly negative AI indicates almost no firing during onset. The AI for each deflection direction of Cell 2 is shown in Figure 3E, and aggregated for all neurons and directions in Figure 3G. Cells are ordered by the mean value of AI across all directions. Note that cells which spike preferentially at onset (top, red cells, “RA-like”) transition smoothly into cells that spike less during onset than average (bottom, blue cells; “SA-like”). This smooth transition is maintained if only the preferred direction (PD) of each neuron is considered (Figure 3G, right column).

Consistent with previous studies^33^, Vg adaptation properties often depend strongly on deflection direction; some cells exhibit positive AI for some deflection directions and negative AI for others. Previous work classified each neuron into an “adaptation category” based on the number of directions in which it responded in an RA-like manner^33^. Similarly, we defined a neuron to be RA-like for a given direction if the AI for that direction is positive. The number of directions for which a neuron responds in this RA-like manner is termed its “adaptation category”; cells are evenly distributed across adaptation category (Fig. 3F).

### Neural responses are correlated with many stimulus components, but these stimulus components are tightly correlated

The 3D whisker shape and rotation measured from video can be used to model the mechanical signals at the whisker base. Decomposing the forces and moments into their x, y, and z components yields six quantities, while the two angular rotations, Δ*θ* and Δ*ϕ*, are proxies for the forces associated with the follicle rotating within the tissue. These eight quantities and their derivatives form a total of 16 dimensions that completely describe the whisker’s mechanical state. A tuning map from this 16-dimensional input space onto the average firing rate of each neuron quantifies the neural response during contact. The full tuning map for each neuron can be projected onto each individual input dimension; some of these one-dimensional tuning maps (tuning curves) are shown in Figure 4A for an example cell, and additional two-dimensional tuning maps are shown in Figure 4B. The total number of spikes in these histograms is 11,010; bins in which fewer than 10 observations occurred were omitted from the map.

**Figure 4:**
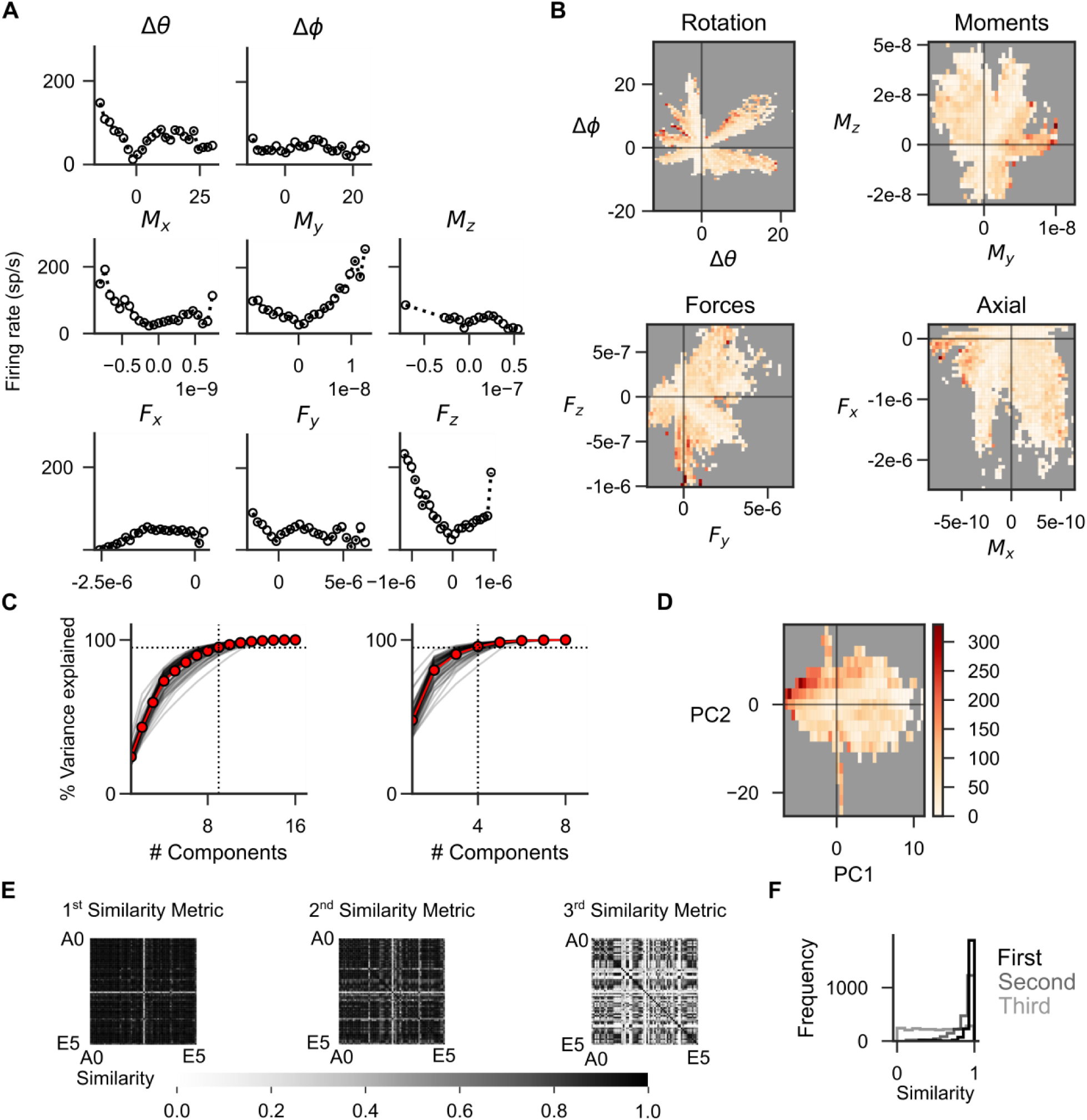
State spaces for each whisker fall in a lower dimensional subspace. In A,B, and D, the number of spikes comprising the histograms is 11,010. **(A)** One-dimensional tuning maps: firing rate as a function of individual components of bending and rotation for an example cell. Units are degrees (top), Newton-meters (middle), and Newtons (bottom). **(B)** Two-dimensional tuning maps: firing rate as a function of pairs of components for the same neuron as in **A**. Color indicates firing rate, scale as in **D**, units as in **A**. Bins in which fewer than 10 observations occurred were omitted from the map. **(C)** The cumulative percent variance explained by subsequent additions of ordered principal components for all whiskers (gray) and on average (red) when derivatives are included (left) and excluded (right). A 95 ms window LOESS smoothing was used before derivatives were computed. Dashed horizontal lines indicate 95% variance explained threshold; vertical lines indicate number of components needed to exceed the 95% threshold. Without the derivatives, three components are required to explain 95% of the variance in the 8-dimensional space; with derivatives, eight components are required to account for 95% of the variance in the 16-dimensional space. **(D)** Two-dimensional tuning map of the neuron in **B** for the two first principal components of its input space, which includes derivatives. **(E)** Pairwise comparisons of the similarity between the 3-dimensional subspaces spanned by the first 3 PCs for all whiskers. Axes are ordered by whisker identity (A0(*α*)-E5). Multiple whiskers of the same row and column identity were included because whisker geometry differs across rats and stimulation pattern differs across trials. Shading is the value of *S*_*i*_ = cos (*θ*_*i*_). **(F)** Histogram of the number of pairwise comparisons with a given value of cos(*θ*_*i*_).

Generally, tuning curves for all cells showed structure in most of the one-dimensional projections: the firing rates of most Vg neurons correlate with most individual parameters. However, these correlations could result from intrinsic covariation between individual stimulus components – for instance, *M*_*y*_ covaries with *dF*_*z*_, where *d* is the distance from the whisker base to the contact point. To quantify these covariations, we performed PCA on the input space for each whisker (with and without derivatives); the cumulative percent variance explained is shown in Figure 4C for all whiskers.

Instead of using the physical quantities as independent variables, tuning maps can also be obtained as a function of the principal components (PCs) of the input space. Example low-dimensional tuning maps for the two first PCs are shown in Figure 4D for the same neuron as in Figure 4B. Derivatives are included in the PCA decomposition; notably, PCA eigenvectors tend to represent combinations of either physical quantities or their derivatives, and rarely mix both (Supplementary Fig. 3).

Because all physical quantities are measured independently of the whiskers’ orientation on the head^36^, we can quantify the similarity between the leading low-dimensional PC representations of stimuli space across different whiskers. This analysis is equivalent to asking whether different whiskers constrain the physical input spaces in a similar way. To quantify the similarity between PC representations, we used a similarity metric (*S*_*i*_) that generalizes the dot product to measure the angles between two subspaces rather than the angle between two lines (see *Methods*). This approach is known in formal mathematics as “canonical angles analysis” (*S*_*i*_ = cos (*θ*_*i*_))^37^, and resembles the “subspace overlap” used in previous work ^38^. Lines are one-dimensional; there is only one angle between two lines. In general, there are as many angles, or similarity metrics *S*_*i*_, as dimensions in the subspaces. A value of *S*_*i*_ = 1 corresponds to two parallel lines, one within each subspace, indicating high similarity between the spaces. A value of *S*_*i*_ = 0 corresponds to orthogonal directions. Multiple angles between subspaces are ordered by decreasing *S*_*i*_ (i.e., increasing *θ*_*i*_).

We quantified input space similarity across whiskers by computing *S*_*i*_, *i* = 1,2,3 for pairwise comparisons between all stimulated whiskers, based on the subspaces spanned by the first three PCs for each whisker (Fig. 4E). Because the applied stimulation varied with each experiment, and because the mechanical properties of each whisker are unique, the observed mechanical stimulus space is also unique, even for experiments on whiskers that have the same row and column identity. Most pairwise comparisons were found to be very similar: *S*_*i*_ > 0.82 > 0.54 > 0.05 for 95% of the three leading angles; weak clustering occurs only for the third canonical angle. This result implies substantial similarity of the relevant input space across whiskers.

### GLMs reveal encoding of rotation and distributed tiling of explored stimulus space

The preceding sections have shown that Vg neurons encode multiple stimulus features, and that stimulus features themselves are strongly correlated. The one- and two-dimensional tuning maps of Figure 4A,B provide intuition for the neural representation of select stimulus features, but fall short of describing the full neural response to the presented stimuli. A full description would require knowing the average firing rate in response to any arbitrary point in the stimulus space, and thus fitting a tuning histogram such as those in Figure 4A to the full 16-dimensional stimulus space. This goal cannot be achieved by systematic and exhaustive exploration, and requires a modelling approach.

We therefore implemented a recent formulation of Generalized Linear Models (GLMs)^25^ that allows for multiple input filters, and thus for modeled neurons to be excited by inputs in multiple directions within the 16-dimensional space. All models were fit with 3 filters, each defined in the 16-dimensional input space (see *Methods* and Supplementary Fig. 4). Only stimulus values at the current time bin were accessible to the models, which included no input history. A parametric nonlinearity (5 parameters) was fit for each filter, bringing the number of model parameters to [(16 + 5) ∗ 3] = 63 per neuron; overfitting was minimized via 10-fold cross-validation.

As illustrated for the example neuron in Figure 5A, we used the models to predict the spike rate with millisecond resolution, smoothed the observed spike train with a Gaussian kernel whose standard deviation *σ* was varied exponentially from 2 to 512 ms, and then compared the predicted rate to the smoothed rate for each value of *σ*. The Pearson Correlations comparing the observed and predicted rates are shown in Figure 5B; correlations are calculated only during contact periods. On average, models best predicted the observed spike rate for *σ*=32 ms, but for many neurons high prediction accuracies are achieved for *σ* as short as 2 ms. Moreover, performances achieved with *σ* = 16 and 8 ms were statistically indistinguishable from that achieved for *σ* = 32 ms (Tukey’s post-hoc test). The relationship shown in Figure 5B is nonmonotonic; model performance drops as the value of *σ* increases, indicating that the models accurately predict high resolution temporal structure in the spike trains rather than fitting the average spike rate. The median correlation value was 0.69 (IQR = [0.55,0.81]) for *σ*=32 ms, with a minimum=0.08 and a maximum=0.91.

**Figure 5:**
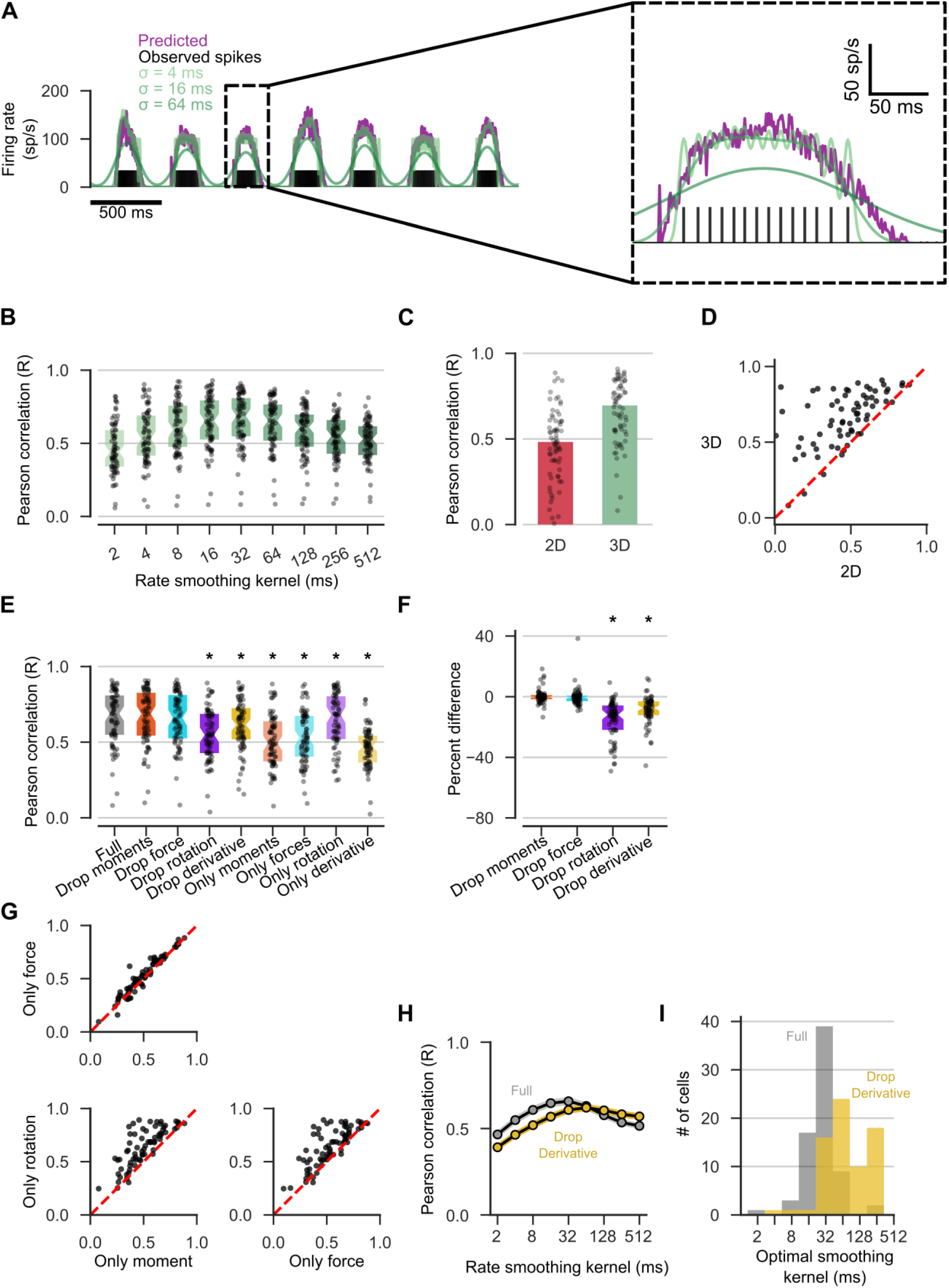
Statistical modeling of Vg neurons. Dots indicate individual models, bars are medians, boxes are median ± 1 quartile unless indicated. For all panels except **B**, Pearson Correlation values are used to compare predicted firing rate to observed rate smoothed with a Gaussian kernel with *σ* = 32 ms. **(A)** Observed spiking (black vertical lines) is converted to estimated rate by smoothing with a Gaussian kernel (green, light to dark: *σ* = [4, 16, 64] ms). Three smoothing resolutions are shown, but nine are computed. Predicted rate is shown in purple. Inset shows a single deflection. **(B)** Correlations for GLM predicted rates compared to the observed spike rates, smoothed with Gaussian kernels with increasing *σ* (light to dark). **(C)** Correlation of models with access to either 2D or 3D physical information. (**D**) Pairwise comparisons of performance for models based on 2D or 3D physical information. **(E)** Performance of models with access to subsets of the full input space. Asterisk indicates significant difference from performance of full model (Wilcoxon signed-rank test, p<0.05). **(F)** Percent difference between the performance of the full model and the model without access to a subset of inputs, (*R*_*subset*_ − *R*_*full*_)/(*R*_*full*_), aggregated over all neurons. **(G)** Comparison of performance of models with access only to inputs of a single class. **(H)** Comparison of performance of models with and without access to derivatives for rates smoothed with increasing values of *σ*. Shown are means ± S.E.M. **(I)** Models with access to derivatives (grey) better predict rates smoothed with lower values of *σ* than those without (yellow).

To determine how much information is gained from the full 3D whisker shape compared to a 2D projection (Fig. 5C,D), the 3D whisker shape was projected into the top camera view. The four mechanical variables associated with contact, {*M, F*_*x*_, *F*_*y*_, Δ*θ*}, were computed using an established 2D model^24,39^. We mapped this 8-dimensional input space (including derivatives) onto the average firing rate of each neuron, and, as in the 3D case, modeled each map using a GLM with three filters. 70 neurons were fit with both 2D and 3D models. We again found that the accuracy of the predicted firing rate prediction was non-monotonic with the value of *σ* used to smooth the spike trains. For *σ*=32 ms, 2D model performance was significantly worse than that of 3D models (Wilcoxon signed-rank test, W=51.0, p<0.001; median=0.48, IQR = [0.37,0.60]). When the 3D model for a given neuron is compared with its 2D counterpart, nearly all models (60/70) perform better with 3D information (Fig. 5D). The median performance increase from 2D to 3D was 29.8% (IQR **=** [13.0%,73.9%]).

In order to determine the relative contribution of each input component to firing rate prediction, we performed a dropout analysis in which models access to information was increasingly restricted (Fig. 5E). For these analyses, correlations were computed using only the *σ*=32 ms smoothed rate. Models without derivative information or rotation information perform significantly worse than the full model (Wilcoxon signed-rank test, W=17, p<0.001 median=0.61, IQR = [0.52, 0.72]; W=209, p<0.001, median=0.55, IQR = [0.43, 0.69], respectively), indicating that these quantities are important for firing rate prediction.

We then asked how well models performed when given access to only one class of inputs: Moments, Forces, Rotations, or Derivatives (Fig. 5E). All models with access to only one class perform significantly worse than the full model (Wilcoxon signed-rank test, p<0.001). Models with only rotation components perform only slightly less accurately than the full models (median=0.68, IQR = [0.52,0.80], W=599), indicating that a majority of the variance in firing rates is accounted for by the rotational components of the input. Models with access to only derivatives perform worst of all dropout models (median=0.47, IQR = [0.36,0.54], W=32), while those with access to just moments or forces perform moderately well (median=0.49, IQR = [0.37,0.64], W=31; median=0.53, IQR = [0.40,0.67], W=24). The percent difference (*R*_*subset*_ − *R*_*full*_)/(*R*_*full*_) quantifies whether the loss of either rotation or derivative information is detrimental on a per neuron basis. Results aggregated over neurons (Fig. 5G) indicate that some neurons are more amenable to modelling than others, and that force and moment both carry important but incomplete information about the response, while rotational variables are the most informative.

Unsurprisingly, models that include derivative information exhibit improved temporal precision. Specifically, models that include derivatives perform better than those that do not, when their predictions are compared to spike rates smoothed with *σ*<64 ms (Wilcoxon signed-rank test, p<0.001, Bonferroni corrected for all kernel sizes tested, Fig. 5H). Moreover, identifying the value of *σ* for which the firing rate of each neuron is best predicted shows that most neurons are best predicted at shorter timescales when derivative information is included (Wilcoxon signed-rank test, n= 74, W=0, p<0.001, Fig. 5I).

Lastly, we analyzed the coefficients that characterize the GLM filters. These coefficients are organized as three vectors in the 16-dimensional input space; these vectors define a 3D subspace that can be interpreted as a complex, three-dimensional “mechanical receptive field” for each neuron. This receptive field cannot be visualized using standard tuning curve analyses, but it provides a more complete description of a neuron’s response properties, and allows us to ask two important questions. First, do the neural representations correspond to low dimensional structure (i.e., the principal components) of the stimulus space itself? And second, how do the complex representations of different neurons compare to each other?

Both of these questions can be addressed by computing again the similarity metric *S*_*i*_ based on canonical angle analysis^37^. This computational approach is schematized in Figure 6. For each neuron, we calculated the similarity between the neural representation and the principal components of the stimulus space. Surprisingly, the neural representations are not strongly similar to the principal components: *S*_1,2,3_ = [0.36 ± 0.13, 0.14 ± 0.07, 0.03 ± 0.03]. Figure 7A shows the similarity metrics for all cells, ordered by decreasing similarity. Corresponding histograms are shown in Figure 7B. Thus, Vg neurons do not seem to preferentially encode high-variance combinations of mechanical stimuli.

**Figure 6:**
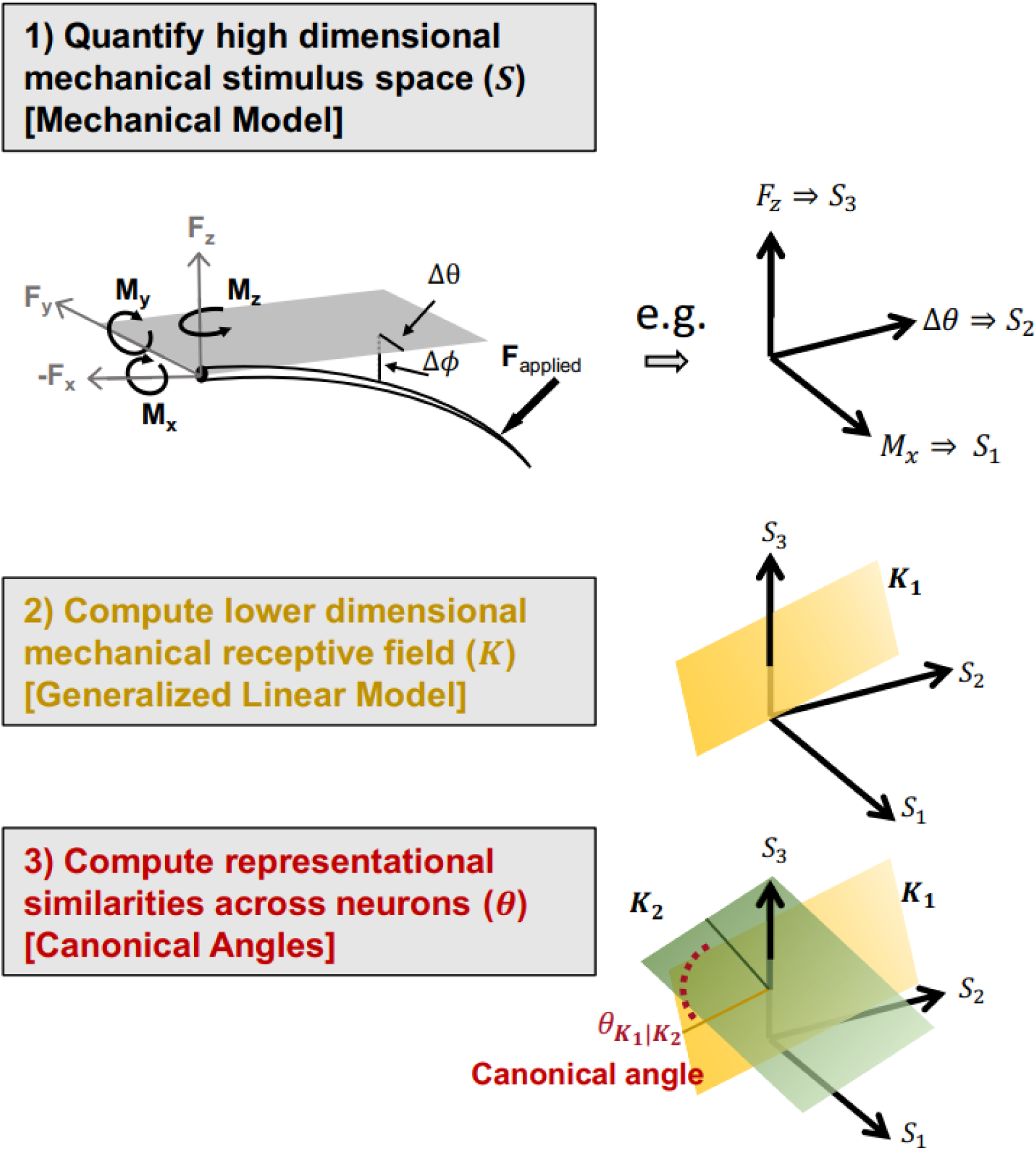
Mathematical methods schematic: An illustration of the mathematical and modelling methods. (1) The 3D whisker shape is extracted from each video frame and viewed from a whisker centered coordinate system. From the 3D whisker shape, we measure the rigid body angular rotations from rest (Δ*θ*, Δ*ϕ*), and estimate the applied force using a 3D mechanical model of whisker bending^24^. The applied force can be decomposed into the component forces at the base of the whisker; knowledge of the contact point allows for the computation of component moments. These mechanical quantities and their derivatives define a 16-dimensional stimulus space (*X*). A toy example stimulus space composed of just three dimensions is shown to the right. (2) We fit low dimensional Generalized Linear Models (GLMs) to match the observed spiking of each neuron. Each GLM model finds a lower dimensional subspace embedded in the higher dimensional, full stimulus space. This subspace, shown here as the plane *K*_1_ (yellow), provides a representation of the stimuli that explain the firing of neuron 1. (3) We compute the similarities between the neural representations of different neurons by calculating the angles between the corresponding neural representational subspaces (*K*_1_, *K*_2_).

**Figure 7:**
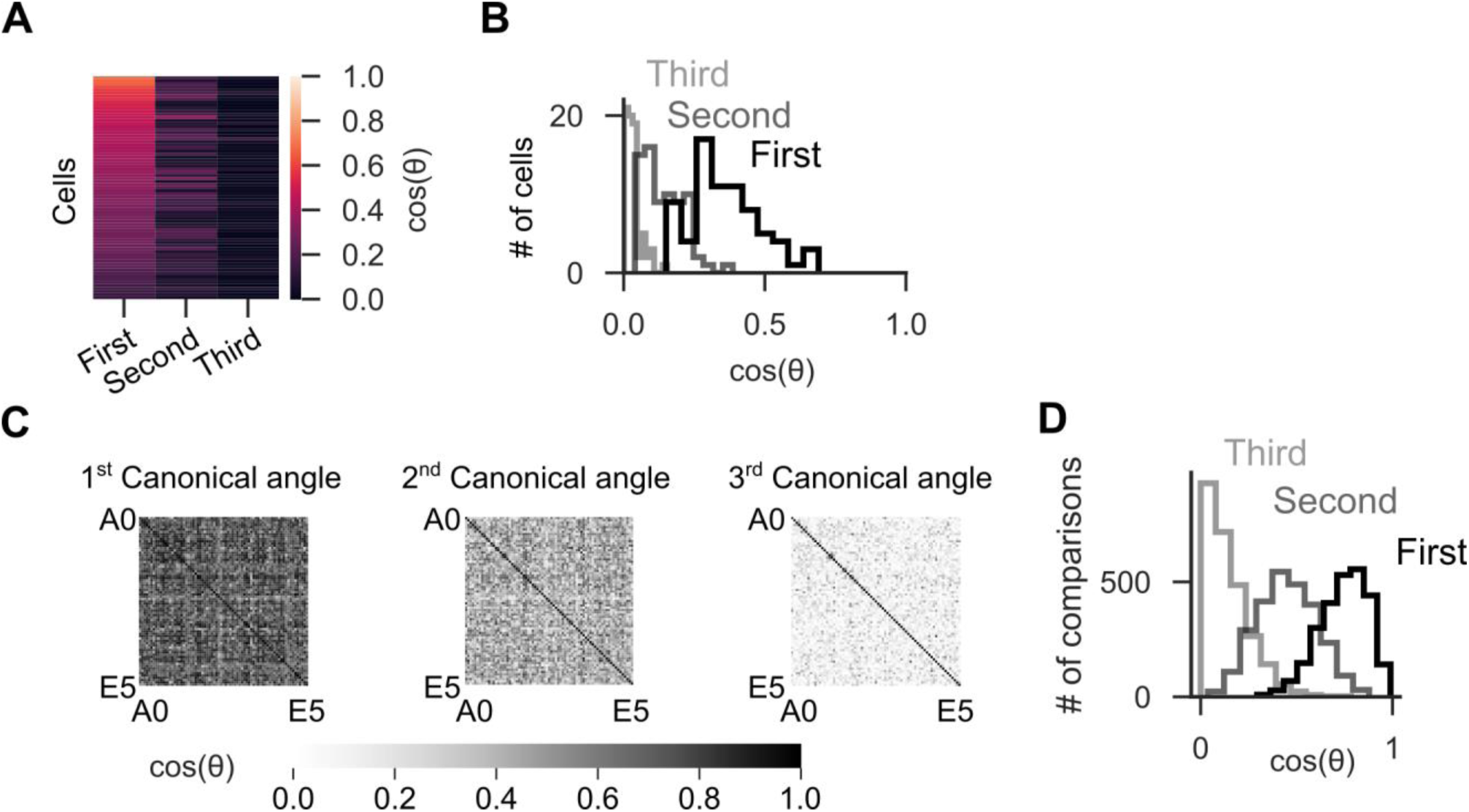
Neural tuning is well distributed across the population and covers the full input space. **(A)** For each neuron, the similarity between the stimulus space principal components and the neural representation is shown. The value of *S*_*i*_ between the input subspace and the neural subspace is color coded (scale on right). Each row is a neuron; neurons are ordered by the value of the first canonical angle. **(B)** Histograms of the values of *S*_*i*_ over all neurons for the three similarity metrics. **(C)** Pairwise similarities of neural representations across all recorded neurons. Axes are ordered by innervated whisker - A0(*α*) to E5. Gray intensity (scale on bottom) indicates the value of *S*_*i*_. **(D)** Histograms of the values of *S*_*i*_ across all pairwise comparisons of the neural subspaces.

We next calculate the similarity between the neural representations themselves to determine if there are identifiable “groups” or “classes” of neurons that share similar representations. The existence of functional cell classes would be indicated by subpopulations of neurons that are similar to other neurons within their class, and dissimilar to those outside their class. Pairwise comparisons are shown in Figure 7C and the resulting histograms in Figure 7D.

In contrast to multiple previous studies that have found distinct Vg cell classes^4,11,19,28,40,41^, the neural representation subspaces overlap only moderately: *S*_1,2,3_ = [0.74 ± 0.12, 0.45 ± 0.15, 0.14 ± 0.11], mean ± S.D., and do not cluster. The lack of clusters and the lack of bimodal distributions in the histograms of the similarity metrics indicate that Vg neurons should not be grouped into classes of cells that represent similar mechanical information.

Thus, the neural representations of Vg neurons do not preferentially cover the low-dimensional representation of input space found by PCA, and their tuning spaces only moderately overlap, but do not cluster. This suggests that as a population, Vg neurons implement a dense, uniform covering of the full input space (Supplementary Fig. 4).

## Discussion

Coding properties of Vg neurons can be fully quantified only if the stimuli employed span the extent of the naturalistic stimulus space. The present work, which allows mechanical stimulus features to be estimated during complex, 3D whisker deflections, shows that individual Vg neurons simultaneously encode multiple stimulus features, that adaptation properties do not categorize into rapidly adapting or slowly adapting groups, and that neural representations overlap and tile the stimulus space. Together, these results suggest a view of Vg coding in which single stimulus features cannot unambiguously be determined by the activity of a single neuron; instead, features are represented across a population, and may be extracted by more central neurons that integrate information across many Vg neurons. This view contrasts with proposed population codes that segregate behaviorally relevant quantities into separate classes^11,28,40,42^.

### Classical Vg tuning curves are “slices” of possible neural responses to complex stimuli

Previous studies used restricted stimulus sets that vary only a few stimulus dimensions while holding others constant^1-3,7,9,10,32^. The Vg responses shown in these earlier studies can be considered “slices” through multidimensional tuning maps that more completely describe responses to complex stimuli.

For example, in agreement with earlier work^9,10,28,33^, our results show that both arclength of contact and deflection direction strongly influence average firing rate. However, we further show that Vg neurons are jointly tuned to both stimulus features, i.e., arclength and direction cannot be disambiguated based on average firing rate (Fig. 2). Here, the extended GLM models attempt to fully describe these more complex tuning maps from which general coding principles can be inferred. For instance, predictive performance of these models decrease when derivatives of mechanical features are omitted as inputs (Fig. 5E), consistent with earlier work showing joint tuning to features corresponding to stimulus “amplitude” and “velocity”^1-3,7,31,32^.

Recently, studies based on 2D analyses of whisker shape have shown that Vg responses are more accurately described in terms of mechanical rather than geometric variables^18,19,29^, but the 2D work omits crucial information about 3D deflection direction, which modulates the components of applied forces. The Vg responses shown in these 2D studies can thus also be considered as “slices” or marginals of responses to the 3D stimulation used here. Although the marginal neural responses to individual features of the stimulus recapture observations from previous work, a more complete picture emerges when responses to more complex stimuli are analyzed without marginalizing to individual stimulus features.

### Vg neural responses tile the mechanical space

Vg responses are tuned to multiple features of the 3D mechanical stimulus space (Fig. 5); statistical models indicate that their activity is primarily driven by the follicle’s rotation in the skin rather than by whisker bending. Predictive performance of the Vg models degraded if information about rotation is omitted, but does not degrade if information about bending is omitted. Remarkably, models that have access only to bending information still perform moderately well, providing evidence for broad, diffuse tuning to mechanical features.

Given the relationship between bending and rotation (Supplementary Fig. 1), it is possible that neurons encode latent mechanical features that subsume bending and rotation. Principal components of the stimulus space represent such latent features. However, the neural representation does not align strongly with the principal components of the stimuli (Fig. 7): neurons do not encode linear combinations of stimulus features along high variance dimensions. Preferential encoding along dimensions that differ from those that characterize the variance structure of the stimuli is consistent with a diffuse, tiled representation of mechanical information.

Although the present work used passive whisker deflections, we expect the neural coverage of the stimulus to remain stable regardless of whether exploration is active or passive. A Vg neuron’s stimulus representation cannot depend on context, as it receives no neuronal inputs. Rather, the statistics of the stimuli may differ in different contexts. In this study, neural responses are likely dominated by rotations of the whisker because the muscles holding the whiskers are relaxed. During active whisking, the muscles contract around the follicle, resisting passive rotation within the skin and cause the whisker to bend rather than rotate^18,43^. The resulting differences at the base of the whisker will correspondingly alter the effective stimulus space for the awake animal such that bending is more prominent during active whisking. However, the neural representation of the mechanical space itself ought to remain unchanged, with the caveat that follicle stiffness may change in the awake animal. More central neurons would therefore extract features of the stimulus from a tiled and distributed representation in the Vg population, while accounting for the invariance of the map from stimulus space to Vg neural activity across passive and active contexts.

### Adaptation characteristics of Vg neurons lie along a continuum

Vg neurons are typically classified as rapidly and slowly adapting (RA/SA). This classification is conceptually intuitive, simplifies analyses, and is consistent with the existence of genetically and physiologically distinct mechanoreceptor types^44-46^. However, during naturalistic stimulation we observe a continuous distribution of adaptation properties that depend on direction (Fig. 3). This finding supports earlier interpretations of non-categorical adaptation responses^12,32,33^.

One interpretation of this finding is that Vg neurons respond both to mechanical features and their temporal derivatives, with differential weights, such that some neurons respond very little to derivatives (SA-like), and others exclusively so (RA-like). The precise balance of these weights is likely affected by various aspects of follicle configuration: the physiological class of the mechanoreceptors, the arrangement and location of the mechanoreceptors in the follicle, and the tissue dynamics of the follicle/mystacial pad^43,47^. The diversity within and across classes of mechanoreceptors likely generates diverse responses in the Vg population, resulting in a more complete tiling of the possible stimulus space, and thus avoiding gaps in the information conveyed to more central neurons^48,49^.

### On the plausibility of a dense code

To fully describe the trigeminal population code would require simultaneous recordings from many Vg neurons. In this study, Vg neurons were individually and sequentially recorded from different animals. Nonetheless, several lines of reasoning suggest that the most parsimonious interpretation of the present results is that the population of Vg neurons represents the stimulus space via a dense code.

First, each Vg neuron responds to many different mechanical states of the whisker (Fig. 5) and to many stimulus features (Fig. 2); many Vg neurons are thus required to fully represent any given stimulus. Second, Vg coding properties are continuously distributed across all recorded neurons (Fig. 3 and 7). Direction selectivity index, temporal adaptation patterns, alignment of the neural representation with the stimulus principal components, and alignment between neural representations, all vary smoothly across the entire population of recorded neurons, indicating a tiling of the input space. Finally, Vg neurons exhibit a wide range of firing rates (Fig. 2,^12,16,32^), consistent with a dense coding scheme^50^.

Vg neurons must represent a large range of mechanical stimuli in multiple behavioral contexts, including active and passive touch, texture discrimination, collisions with objects, non-contact whisking, and airflow exploration^19,51-53^. A dense coding scheme such as the one proposed here would offer several distinct advantages. It offers robustness against noise in individual neurons, and even their loss. It has a high representational capacity, a useful property given that there are only about 200 to 300 Vg neurons per whisker^54^. A distributed, dense code would allow for individual Vg neurons to be informative of stimuli under many contexts, without filtering out information at this early stage. In this way, the Vg population could represent arbitrary stimuli in the space of all possible stimuli, and allow more central neurons to extract those features that are relevant in the context of the animal’s ongoing behavior and motor actions.

## Methods

All procedures involving animals were approved by the Northwestern Animal Care and Use Committee. A total of 22 female Long Evans rats between 3 – 6 months were used.

### Surgical procedures and electrophysiological recordings

Animals were anesthetized with a ketamine-xylazine-acepromazine cocktail administered intraperitoneally (60mg/kg ketamine, 3.0 mg/kg xylazine, 0.6 mg/kg acepromazine). After deep anesthesia was induced, the fur from the left whisker array was removed with depilatory cream (Nair) to increase contrast near the proximal region of the whisker close to the basepoint. Care was taken to minimize contact between Nair and the whiskers, and to wash off the Nair as soon as possible with saline. If the shape of a whisker was visibly altered by the fur removal procedures, it was removed from the array prior to recordings.

The head was immobilized with ear-bars to a custom stereotaxic device, and three stainless steel skull screws were inserted on the dorsal aspect of the cranium. Prior to the surgery, a non-insulated silver wire had been soldered to one of the skull screws to serve as a ground wire for electrophysiological recordings.

An approximately 1 mm diameter craniotomy was made over the left hemisphere, 2 mm caudal to bregma and 2 mm lateral to the midline. The skull was leveled to ensure that the bregma-lambda plane was horizontal, and a dental cement (methyl methacrylate) “bridge” was formed to connect the skull screws to the right side of the stereotaxic device. This procedure allowed for removal of the bite support and left ear bar while maintaining a level head position, giving free access to the left whisker array for stimulation. Once the dental cement bridge had set, a single tungsten electrode (FHC 1-3 MΩ) was centered over the craniotomy and lowered to a depth of ∼10 mm, until whisker responsive field potentials could be heard in audio monitoring of the amplified electrode signal during manual stimulation of the entire whisker array. We then waited ∼5-10 minutes to allow the brain to relax after the initial penetration before advancing slowly to isolate a unit that responded only to the deflection of a single whisker.

Once a single unit was isolated, the whisker associated with that neuron was visually isolated to ensure high contrast in both front and top camera views. A white paper background was placed behind the whisker to provide a uniform background for robust tracking in the front camera. Surrounding whiskers were either trimmed or placed carefully behind the paper background. Care was taken not to deform the whisker of interest or the surrounding mystacial pad.

A custom LED sheet with a transparent white plexiglass diffuser was used as the background lighting for the top camera. An adjustable Neewer CN-160 LED array was used as foreground lighting in the front camera field of view. High speed video from two identical top and front cameras was recorded directly at either 300 fps (Teledyne Dalsa HM640) or 500 fps (Mikrotron 4CXP) using StreamPix 7. Front and top cameras were synchronized by way of clocked 5V TTL to initiate exposure of each frame in both cameras from the same source. At the end of each experiment we recorded images of a checkerboard pattern with 2 mm squares in the field of view of both cameras; these images were later used for camera calibration and for calculation of the 3D whisker shape.

Neural signals were amplified using a A-M systems 4 channel amplifier, with a 10 Hz to 10 kHz hardware filter, at 1000x gain. Amplified signals were acquired via Measurements Computing DT304 card using Datawave SciWorks v8. After acquisition, signals were digitally bandpass filtered at 300-6000 Hz before spike sorting with KlustaKwik^55^.

During recording, whiskers were manually deflected with a graphite probe (0.3 mm diameter) in 8 cardinal directions with respect to the emerging axis of the whisker. Deflections were applied at 2-3 distances along the whisker (arclengths), and at approximately 2 speeds, for a total of approximately 32-48 different categories of deflection. Each category of deflection was repeated ∼20 times for each whisker. Care was taken to minimize slip along the length of the whisker during a deflection. Neural signals and subsequent stimulus quantifications were analyzed using custom python and MATLAB code based on the neo and elephant python packages.

### 3D whisker reconstruction

Whiskers were first tracked in 2D automatically using the software “Whisk”^30^. All tracked videos were manually inspected to verify that the desired whisker was adequately tracked in each frame and view. Videos in which the whisker was not adequately tracked (e.g., background edges were labeled as the whisker, manipulator was tracked as the whisker, tracking did not extend sufficiently to the tip or to the base) were omitted from further analyses. In order to reconstruct the 3D whisker, the basepoint position needed to be accurate in both front and top images. Since tracking of the basepoint location was occasionally noisy or unreliable, particularly in the front view, a mask outlining the rat head was manually created for each video and view. Any part of the tracked whisker that fell within the mask was then removed. The distance between the edge of the mask and the basepoint was calculated, and the whisker was linearly extrapolated back to the basepoint. This gave an accurate and temporally smooth re-creation of the base segment and basepoint of the whisker in each view.

The two 2D tracked whisker shapes (one from each camera view) were cleaned by first removing and interpolating over mis-tracked basepoints via a median filter (window = 5 frames) and outlier deletion (Grubbs test *α* < 10^−8^). Next, the entire 2D whisker shape in each frame was smoothed with a spatial LOWESS filter (span = 15% of whisker length).

In order to create 3D reconstructions of the whisker, the two cameras had to be first calibrated. This involved computing the intrinsic properties of each camera (focal length, principal point, distortion, and skew), as well as the relationship of the cameras to each other (rotation and translation). These procedures were done with the Caltech Camera Calibration Toolbox, OpenCV, and custom MATLAB and python code.

Once the cameras were calibrated, it became possible to calculate the location of an arbitrary 3D point in an external reference frame based only on two “corresponding” points in the two 2D camera views. In the case of the tracked whiskers, however, the only two available corresponding points are the base and the tip of the whisker, as there are no features on the whisker itself that can be identified as the same point in both camera views.

Therefore, in order to reconstruct the full shape of the whisker, we used an iterative optimization to find the best 3D whisker shape that minimized the back-projection error, where back-projection refers to the 2D projection of the estimated 3D whisker onto either camera. The back-projection error is simply the Euclidean distance between the back-projected whisker and the actual, imaged whisker, summed over all back-projected points. The basepoint is chosen as an initial corresponding point. We then randomly sample a 3D point at some distance *s* from the basepoint in a random direction, and compute the back-projection error for that point. We continued to sample random 3D points at the same distance *s* from the basepoint to find the point with the minimal back-projection error. This optimal point became the next tracked point along the 3D whisker, and the origin for the next search over random 3D points at a distance *s*. Subsequent 3D points were added in the same manner, until either the cumulative back-projection error exceeded a preset threshold, or the whisker began to fold back on itself. The latter happened when the next optimal point was behind the previously fit point; this indicated that the whisker had been completely tracked and the folded point was then discarded. This process was carried out for each tracked frame for all videos. Quality of fits were inspected manually by observing the shape of the 3D whisker over time, viewing the overlap of the 2D back-projections with the original tracked 2D whiskers, and monitoring the temporal trajectories of the base and tip points for large deviations.

### 3D contact point estimation

We needed to calculate the 3D point of contact of between whisker and manipulandum, both to quantify the arclength at which contact occurred, and because the contact point is needed to compute the applied forces and moments in the 3D mechanical model^24,39^. A difficulty was that the manipulator had no corresponding points, making the 3D reconstruction of the manipulator impossible. Instead, the manipulator was tracked as a 2D line in each frame via a custom written spatially and temporally constrained Hough line search. We then computed the intersection of that 2D manipulator line with the previously computed 2D back-projection of the 3D whisker for the corresponding frame and camera view. This resulted in a 2D point in the camera view. Since the 2D back-projected whisker consisted of the same number of tracked points as the 3D whisker, the node of the 2D back-projected whisker that was closest to the intersection point was designated as the 2D contact point. This point corresponded to a point on the 3D tracked whisker, which was then deemed the 3D contact point.

This approach had the advantage that the manipulator needed to be tracked in only one camera view per frame, and that the 3D contact point was constrained to fall exactly on the reconstructed 3D whisker. The latter would not be guaranteed if the manipulator was reconstructed in 3D in the same way as the whisker. Moreover, it was likely that for any deflection in any given direction, the manipulator was relatively perpendicular to the field of view in one camera, and so line detection was robust. Accurate tracking of the manipulator in all frames was ensured in two ways: first the custom written tracking software would warn the user if large spatial or temporal changes were detected, which prevented errant tracking in almost all cases; second, tracking quality was monitored online in all videos during tracking and confirmed as acceptable by a second user offline.

### Contact determination

Because stimulation was manually delivered, the actual time of contact onset was not known, and there was no temporally repeatable stimulus onset. To determine contact manually, the most accurate method would be to observe the recorded video and determine the first frame in which the tip of the whisker moved significantly. It was infeasible to do this for every contact and cell recorded, so we extracted the tip position of the whisker from the two camera views, resulting in a 4D temporal trace of tip position coordinates (Top (x,y), Front (x,y)). Naïve techniques in which contact was assumed to occur when these traces crossed a threshold were not robust enough to adequately distinguish contact from non-contact. Instead, we trained a temporal convolutional neural network to distinguish between contact and non-contact for every frame. We first whitened the 4D tip position within a video to have zero mean and isotropic unit variance. We then labeled a random selection of ∼1 million frames (amounting to approximately 10% of all data) across all experiments as either contact or non-contact. Labeling was achieved by viewing the randomly sampled segments of the 4D temporal trace of the tip position. A user visually determined the onset and offset of a contact period, marked by a significant deviation in the tip position. Frames between onset and offset were labeled as contact, and the remaining frames in the segment were labeled as non-contact. We split the labeled data into a training and a test set, and chose the architecture of the network to be able to correctly classify 97.7% of the training set and 97.4% of the test set. We then applied the trained neural network to the remaining ∼9 million frames to get a contact/non-contact label for every frame, and manually verified the contact predictions by observing the labeled 4D temporal traces for all frames and correcting the predicted time of contact as needed. Ultimately, every contact was inspected manually based on the 4D temporal traces of the whisker tip, but the neural network drastically reduced the effort that would have been needed to manually label every one of the ∼10 million recorded frames.

### 3D mechanical models

The mechanical models used here to calculate the three components of force and three components of moment at the base of the whisker have been described previously^56^. All calculations were done in whisker-centered coordinates, in which the whisker basepoint is centered at the origin, and the whisker is rotated such that the approximately linear portion of the base segment of the whisker is colinear with the x-axis and the initial curvature of the whisker lies in the x-y plane. Mechanical models take the 3D shape of the whisker in the frame prior to each contact onset as the reference whisker for that contact. In each subsequent contact frame during which the whisker is deformed, we estimated the forces and moments required to deform the reference whisker into the whisker shape observed during contact.

As described in previous studies^24,39,56^, the mechanical model approximates the whisker as a tapered, truncated beam. The three components of force and three components of moment {*F*_*x*_, *F*_*y*_, *F*_*z*_, *M*_*x*_, *M*_*y*_, *M*_*z*_} (Supplementary Fig. 1) were computed at the base of each whisker. Importantly, these computations were performed in whisker-centered coordinates for each frame, so that the applied force takes into account only the change in shape of the whisker (i.e., bending). To calculate the rotational component during whisker deflection, we computed the rotation (*θ, ϕ*) required to move the whisker from the camera-centered reference frame to the whisker-centered reference frame at every point in time. The rotation magnitude in each frame was then computed as the change in these angles (Δ*θ*, Δ*ϕ*) from the position of the whisker in the frame prior to contact. Marginal distributions of mechanical and kinematic variables are shown across all contacts and whiskers in Supplementary Fig. 2.

In some cases, we used two additional scalar quantities: the magnitude of the bending moment *M*_*B*_ and the rotation magnitude *M*_*R*_, which quantifies the arc swept in the direction of rotation:

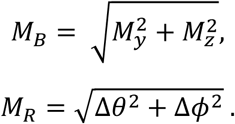

### 2D mechanical models

In order to assess the amount of information gain when moving from 2D to 3D, we calculated the mechanics due to the bending and rotation of the whisker as if we only had information from the top camera. Since the reconstruction of the 3D whisker is an estimation of the 3D shape, it was inappropriate to simply compare the 3D information with the information obtained from direct 2D tracking from the top camera, as the latter is likely more accurate. Instead, we back-projected the estimated 3D whisker onto the top camera, to get a 2D image of the whisker of comparable quality to that of the 3D reconstruction. The contact point was the same node along the whisker as identified during the 3D analysis and did not need to be recomputed. We used the back-projected 2D whisker shape to calculate the angular rotation Δ*θ* as for the 3D models, but now restricted to the 2D projection. We then applied a previously described 2D mechanical model^57,58^, analogous to the 3D model already discussed, to calculate the bending magnitude *M*, the axial force *F*_*x*_ directed into the follicle, and the lateral force *F*_*y*_. The derivatives of these physical quantities were calculated as described for the 3D mechanical quantities. The resulting 8-dimensional stimulus space included {*M, F*_*x*_, *F*_*y*_, Δ*θ*} and their respective derivatives.

### Smoothing, alignment, and upsampling

Errors inevitably occurred at various points in the data analysis: when tracking the 2D whisker shape, when reconstructing the 3D whisker, when calculating the 3D contact point or the force applied to the whisker. These errors caused fluctuations in the calculation of the quantities that characterize the stimulus space. Of particular difficulty were temporal outliers in which data from a given single frame differed greatly from data from the surrounding frames. These outliers added erroneous temporal structure to the various quantities. To deal with this problem, we applied mean absolute deviation (MAD) filtering to remove points that deviated from the mean by more than 10 times the mean deviation, and then applied a Hampel filter, a thresholded median filter with a window size of 5 frames and a threshold of 3 standard deviations^59^. Lastly, we performed Savitzky-Golay filtering on the data, with a span of 7 frames and degree 2, to smooth over high-frequency noise.

Neural data was recorded at 40 kHz, and high-speed video was recorded at either 300 or 500 frames per second. In addition to the voltage from the extracellular electrode, the neural data acquisition system recorded a 5V signal from the top camera that was high for 100 μs at the start of the exposure for that frame. This allowed us to match each video frame to a sample in the neural data. Variables associated with the video images and tracked whiskers (e.g., mechanics, rotation) were linearly interpolated so as to be up-sampled to 1 kHz; spike times were rounded up to the nearest millisecond.

### Contact removal

Due to occasional errors in tracking of the 3D whisker or contact point, there were sometimes entire contact periods that did not meet quality control and had to be removed. To deal with this problem we used the 3 components of moment to detect unacceptable contacts. First, gaps in moment information shorter than 10 consecutive frames were linearly interpolated, and a median filter with a window size of three consecutive frames was then applied to smooth over temporal outliers. Each contact interval was then given a score:

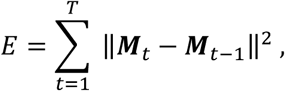

where ***M*** is the 3D moment vector, and *T* is the duration of the contact. If a contact interval had an *E* value greater than 100 times the median value of *E* across all contacts for that given whisker, then the entire contact interval was discarded. We then manually inspected the moment versus time traces for each video and discarded contacts in which the moment signal was dominated by noise.

### Derivative calculations

Temporal derivative information has been shown previously to be of importance to Vg neurons^2,3,7^. In order to calculate the temporal derivative of a physical quantity such as *M*_*x*_, it is customary to temporally smooth the quantity to reduce the effects of sampling noise or small fluctuations in the calculation of the derivative. This procedure is analogous to performing a low pass filter on the quantity whose derivative is to be calculated. Since these quantities were defined to be exactly zero during non-contact, standard smoothing techniques such as Butterworth filters were inappropriate, in that they would have significantly altered the onset/offset boundaries and resulted in non-zero values during non-contact. Instead, we performed a local linear regression (LOESS) smoothing operation on the physical quantities with a window size of 95 ms. After LOESS smoothing, we performed discrete derivative calculations on each of the smoothed quantities.

### Identification of direction and arclength groups

Although the naturalistic stimulation employed here does not allow for repeatability of trials, the applied stimuli could be categorized. Stimulations were applied in 8 cardinal directions relative to the whisker axis, and at 2 to 3 distances from the base (arclengths). In order to average across similar deflections, we labeled each deflection as belonging to a particular direction group and a particular arclength group. To categorize deflections into direction groups, we used Δ*θ* and Δ*ϕ* to represent the angular trajectory of the base segment during a deflection. Since deflections had variable durations, we subsampled the trajectories down to 10 time points per trajectory, and represented the trajectory as a point in a 20-dimensional space of coordinates {Δ*θ*_*i*_, Δ*ϕ*_*i*_}, 1 ≤ *i* ≤ 10. We then applied PCA to the set of 20-dimensional points that corresponded to all deflections of a given whisker, and reduced the dimensionality of this space from 20 to 2 by keeping only the two leading principal components. In this 2-dimensional space, each point provided an abstract representation of one specific whisker deflection. Each of these 2-dimensional vectors was then normalized to unit length, to eliminate the influence of deflection amplitude in the angular grouping. We then clustered all these normalized 2-dimensional vectors into 8 groups using a Gaussian Mixture Model unsupervised clustering algorithm^60^. The procedure was implemented separately for each whisker, and the outcome was visually inspected by color coding all angular trajectories for a given whisker according to cluster label, as shown in Figure 1D. Although little overlap among clusters was observed, different direction groups were not always evenly spaced in angular separation, due to the manual nature of the stimulation. To characterize each deflection direction group, we calculated Δ*θ* and Δ*ϕ* at the apex of the deflection for each deflection in the group, and took the mean of these two maximal values to define the characteristic angular direction of that group.

In each contact frame, we calculated the arclength of contact to be the distance along the whisker from its base to the point where contact was made. We observed only minimal slip of the point of contact along the whisker during a deflection (average slip along whisker during contact was ±0.47mm). Given these small fluctuations in the arclength of contact, we used its median value during a deflection to characterize the whole deflection. We then used Gaussian mixture models to cluster the median arclengths into either 2 or 3 groups. Model selection between clustering into 2 or 3 distinct arclength groups was based on the minimization of the corresponding Akaike Information Criterion (AIC)^61^. If three clusters were found, the deflections were labeled as proximal, medial, or distal; if only two were found, the deflections were labeled as proximal or distal.

### Direction Selectivity Index (DSI)

Several analyses in the present work involve modulation by an angular covariate. In order to quantify this angular influence, we calculated the Direction Selectivity Index (DSI)^34^, defined as:

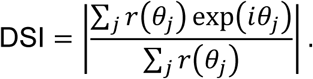

Here *r*(*θ*) is the response variable, typically a firing rate, and *θ*_*j*_ is the value of the angular covariate for the *j*th direction. The DSI is equivalent to 1 − *σ*^2^, where *σ*^2^ is the circular variance of the quantity in question.

### Adaptation Index (AI)

The Adaptation Index (AI) was used to quantify how the firing rate of a given neuron changed over the course of a deflection. We defined the AI as the log of the ratio of the firing rate during the first 10 ms following deflection onset to the average firing rate during the entire deflection:

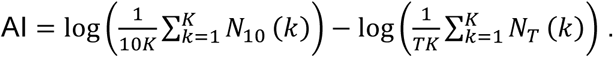

Here *K* is the number of deflections, *N*_10_(*k*) is the number of spikes during the 10 ms following deflection onset for the *k*th deflection, *N*_*T*_(*k*) is the number of spikes during the entire *k*th deflection, and *T* ms is the mean deflection duration, averaged over all deflections of the neuron being characterized by the AI.

### Low-dimensional tuning maps

Similar methods were used to calculate tuning maps in one and two dimensions. In one dimension, the stimulus variable was binned into 25 equal bins; in two dimensions, each of the two stimulus variables was binned into 50 equal bins. The resulting histograms sample the prior probability distribution of the stimulus, marginalized to the corresponding one or two dimensions within the 16-dimensional stimulus space. Bins for which the corresponding stimulus value was observed less than five times were considered empty. For occupied bins, normalized counts were used to estimate the prior probability distribution of the stimuli.

The evoked firing rate of the neuron being mapped was then computed for all occupied bins. The time-dependent spike rate was estimated by convolving the binary spike train with a Gaussian kernel with *σ* = 2 ms. For such small *σ*, conversion to a rate provided smoothing without greatly altering the temporal information. These rates were used to create a new histogram that estimated the expectation value of the firing rate given the stimulus.

To this end, an average spike rate for each stimulus bin was computed over all the times a stimulus value within that bin was observed.

### Generalized linear models (GLMs)

The input space available to a model for predicting the firing response of a specific neuron was the 16-dimensional space consisting of {*M*_*x*_, *M*_*y*_, *M*_*z*_, *F*_*x*_, *F*_*y*_, *F*_*z*_, Δ*θ*, Δ*ϕ*} and the temporal derivatives of these quantities. Each input variable was sampled at 1 ms resolution. The target output for training each neuron specific model was the corresponding binary spike train recorded during the experiment: either a spike was observed (1) or not (0) in each 1 ms bin.

The input *X*(*t*) consists of the values of the 16 stimulus variables at the time *t* of prediction. Since Vg neurons are known to respond to stimulus on fast time scales, sometimes less than 1 ms, and since the temporal resolution of the stimulus is the same as the temporal scale of the Vg response, the model does not need to incorporate a time lag between inputs and outputs or a stimulus history, as has been the case in previous applications^29,62^. The models implemented here were constructed using cylindrical basis functions^25^:

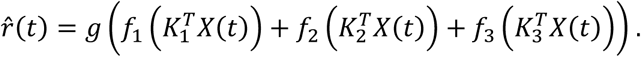

Here *X*(*t*), the stimulus input at time *t*, is projected onto filters *K*_*i*_, 1 ≤ *i* ≤ 3. Each filter is a 16-dimensional vector of weights assigned to each component of *X*; each *f*_*i*_ is a nonlinearity that maps the corresponding projected stimulus into a firing rate. The function *g* is an overall sigmoidal nonlinearity. The functions *f*_*i*_(*x*_*i*_(*t*)), 1 ≤ *i* ≤ 3, each a function of a single scalar 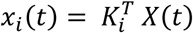, were parametrized as the linear combination of 5 cylindrical basis functions^25^:

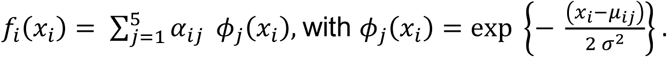

The coefficients {*α*_*ij*_},, 1 ≤ *i* ≤ 3, 1 ≤ *j* ≤ 5, that control the linear combinations of cylindrical basis functions, as well as the additional model parameters {*K*_*i*_}, 1 ≤ *i* ≤ 3, that specify the neural filters, were fit to minimize the negative log-likelihood of the observed spike train given the observed stimulus. All models were 10-fold cross-validated; 90% of the data was used for parameter fitting, and the resulting model was used to predict 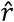 for the remaining 10% of the data. This was repeated 10 times, so that every 1 ms bin for which 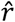 is predicted was at some point not part of the training data used to specify the parameters of the predictive model. Subsequent analyses of the filter weights {*K*_*i*_}, 1 ≤ *i* ≤ 3, for each neuron were performed on mean values obtained by averaging across the 10 cross-validation instances.

For the dropout analysis to establish the relevance of the various input components, we fitted the corresponding predictive models as described above after removing some classes of input components. We found no evidence of overfitting due to too large a parameter space; for instance, the model with the fewest number of parameters (the rotation only model, with only four input components) performed as well as the full model, while other reduced models showed poorer performance than the full model. Models for the 2D whisker description were constructed in the same manner, but based on an 8-dimensional input space that included {*M, F*_*x*_, *F*_*y*_, Δ*θ*} and their temporal derivatives.

We also investigated an alternative approach to modeling the input-output relation of individual neurons, the “spike-triggered mixture model”^26^, based on similar input spaces and employing similar parameters. Results were both qualitatively and quantitatively similar; details about these models and their corresponding results are available on request.

### Pearson Correlations

All models computed a probability of firing 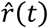 in a given 1 ms bin based on the fitted parameters {*K*_*i*_}, 1 ≤ *i* ≤ 3 for the linear filters and the mixing coefficients {*α*_*ij*_}, 1 ≤ *i* ≤ 3, 1 ≤ *j* ≤ 5 for constructing the fitted nonlinearities *f*_*i*_. This expected rate or firing probability within a 1 ms bin is a continuous variable with values between 0 and 1. The observed spiking *y*(*t*) can be considered as a single observation of the response given an underlying rate *r*(*t*). We computed the underlying rate *r* by smoothing the observed spike train *y* with a Gaussian kernel with standard deviation *σ*. The value of *σ* was varied to investigate the temporal precision of the models. We used *σ* = [2,4,8,16,32,64,128,256,512] ms; this resulted in nine different estimates of *r*, one for each level of smoothing. The Pearson Correlation was then computed between each smoothed *r*(*t*) and the model prediction 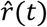. The calculation of correlations was restricted to periods of contact, to avoid overestimation due to silent non-contact periods.

### Similarity metric

The input space considered in the neural response analyses reported here was a 16-dimensional space consisting of the physical quantities {*M*_*x*_, *M*_*y*_, *M*_*z*_, *F*_*x*_, *F*_*y*_, *F*_*z*_, Δ*θ*, Δ*ϕ*} and their temporal derivatives. At any 1 ms time bin during whisker deflection, the stimulus can be represented as a point in this space. For each whisker, the principal components decomposition of the cloud of such points accumulated over many deflections provides eigenvectors and eigenvalues that characterize the stimuli for that whisker, regardless of the response of a Vg neuron associated with that whisker. The eigenvectors are sorted in decreasing order of the eigenvalues, which measure the variance in the corresponding eigenvector direction. Dimensionality reduction to the subspace spanned by the leading eigenvectors maximizes the amount of variance accounted for. Since each whisker has different physical properties (arclength, intrinsic curvature, diameter), the pattern of covariation of the input components will be different for different whiskers, and so will be the resulting eigenvectors.

We have kept the three leading eigenvectors for each whisker, effectively reducing the 16-dimensional input space to a 3-dimensional subspace that is whisker specific. We then asked how similarly oriented were the subspaces associated with different whiskers. The cosine of the canonical angles^37^ between two subspaces quantify this similarity. Given two 3-dimensional subspaces A and B embedded in the full 16-dimensional input space, we considered their spanning vectors *A* = [*a*_1_, *a*_2_, *a*_3_], *B* = [*b*_1_, *b*_2_, *b*_3_]. Here {*a*_*i*_} and {*b*_*i*_} are the three leading eigenvectors that span each subspace; each of them a 16-dimensional column vector. The canonical angles {*θ*_*i*_}, 1 ≤ *i* ≤ 3, follow from

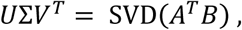

where the diagonal elements of the matrix ∑ are the cosines of the canonical angles,

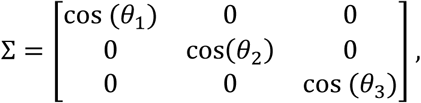

with cos (*θ*_1_) ≥ cos (*θ*_2_) ≥ cos (*θ*_3_), or *θ*_1_ ≤ *θ*_2_ ≤ *θ*_3_.

In addition to comparing input stimulus subspaces across whiskers, we used principal angles to relate the 3-dimensional input subspace associated with a given whisker to the 3-dimensional subspace that best predicted the response of a Vg neuron associated with that whisker. The GLM model for each neuron identified three vectors {*K*_*i*_}, 1 ≤ *i* ≤ 3, each of them a vector in the 16-dimesional input space, and each associated with a preferred input direction for maximal neural response. The orthonormalization of these three vectors provided a basis for a 3-dimensional input subspace that accounted for preferred neural responses. As above, the canonical angles allowed us to compare this 3-dimensional subspace of neural responses to the 3-dimensional subspace that accounted for most of the input variance to the corresponding whiskers.

### Participation ratios

The participation ratio quantifies how evenly distributed the components of a vector are. Given a *d*-dimensional vector *A* = [*a*_1_, *a*_2_ … *a*_*d*_], the participation ratio is defined as:

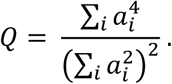

When all the components of *A* are equal, *a*_*i*_ = *a* for all 1 ≤ *i* ≤ *d, Q* attains a minimum value of 1/*d*. When only one component of *A* is non-zero, *a*_*j*_ = *a* and *a*_*i*_ = 0 for all *i* ≠ *j, Q* attains a maximum value of 1. Intermediate values of *Q* quantify the degree of inhomogeneity among the components of the vector *A*.

## Data availability

Data will be made available upon request.

## Code availability

Analyses were performed in MATLAB and python using custom modules and scripts. This software is available upon request.

## Acknowledgements

This work was supported by NIH grants R01-NS093585 to M.J.Z.H. and S.A.S., and F31-NS092335 to N.E.B. We thank Admir Resulaj for technical assistance.

