## Supplemental Figures for "Continuous, multidimensional coding of 3D complex tactile stimuli by primary sensory neurons of the vibrissal system"

### Supplementary Information

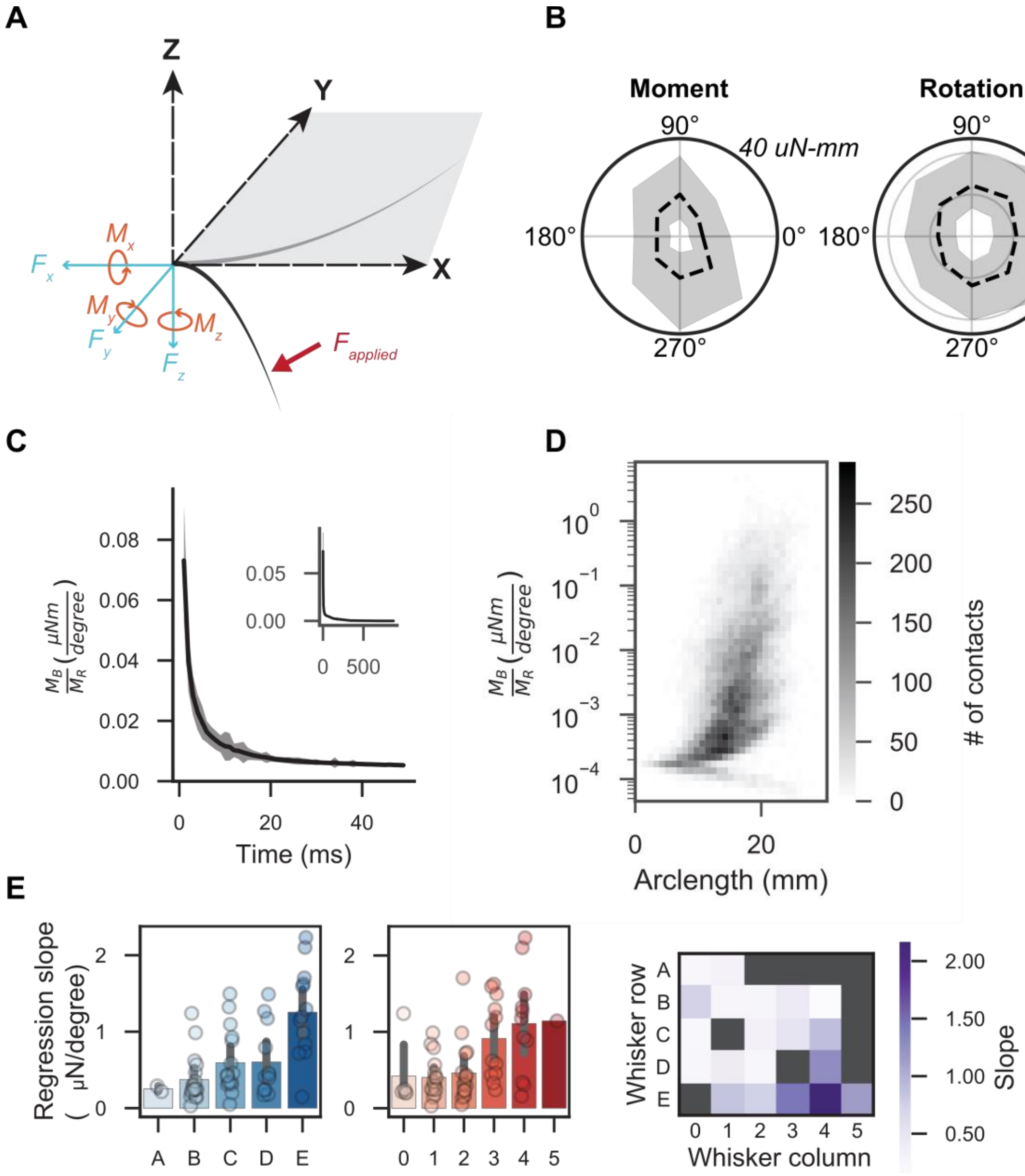


**Supplementary Figure 1: Relationship between bending magnitude and rotation magnitude. (A)** Schematic of the mechanical components of bending in whisker centered coordinates. The base of the whisker at rest (light grey) defines the origin. The x-axis is defined as colinear with the base segment of the whisker, the y-axis as orthogonal to the x-axis in the direction of whisker curvature, the z-axis is orthogonal to the x-y plane. A force applied to the whisker (red) bends the whisker (dark grey). The resulting components of force (cyan) and moment (orange), experienced at the base of the whisker, are determined by the mechanical properties of the whisker and the force applied. **(B)** Median +/- IQR of the magnitude of the bending moment (left) and magnitude of rotation (right) as a function of deflection direction, across all deflections and whiskers. Italic label indicates radial scale. **(C)** Bending precedes rigid rotation of the whisker. The ratio of the magnitude of the bending moment ($M_B$) to the magnitude of rotation ($M_R$) is shown as a function of time after contact onset. Shaded region is 10 x S.E.M. Inset shows the same relationship expanded in time. Data averaged across all contacts and whiskers. **(D)** Ratio of $M_B$ to $M_R$ as a function of arclength of contact, for every contact. Arclength and bending-rotation ratio are median values during a contact. **(E)** Slope of the linear regression of bending-rotation ratio against the arclength of contact for all contacts for each whisker. The relationship between bending and rotation changes as a function of whisker identity. The slope of the linear regression is shown as a function of row identity (left), column identity (middle), and the whisker identity (right). Error bars are 95% confidence intervals.

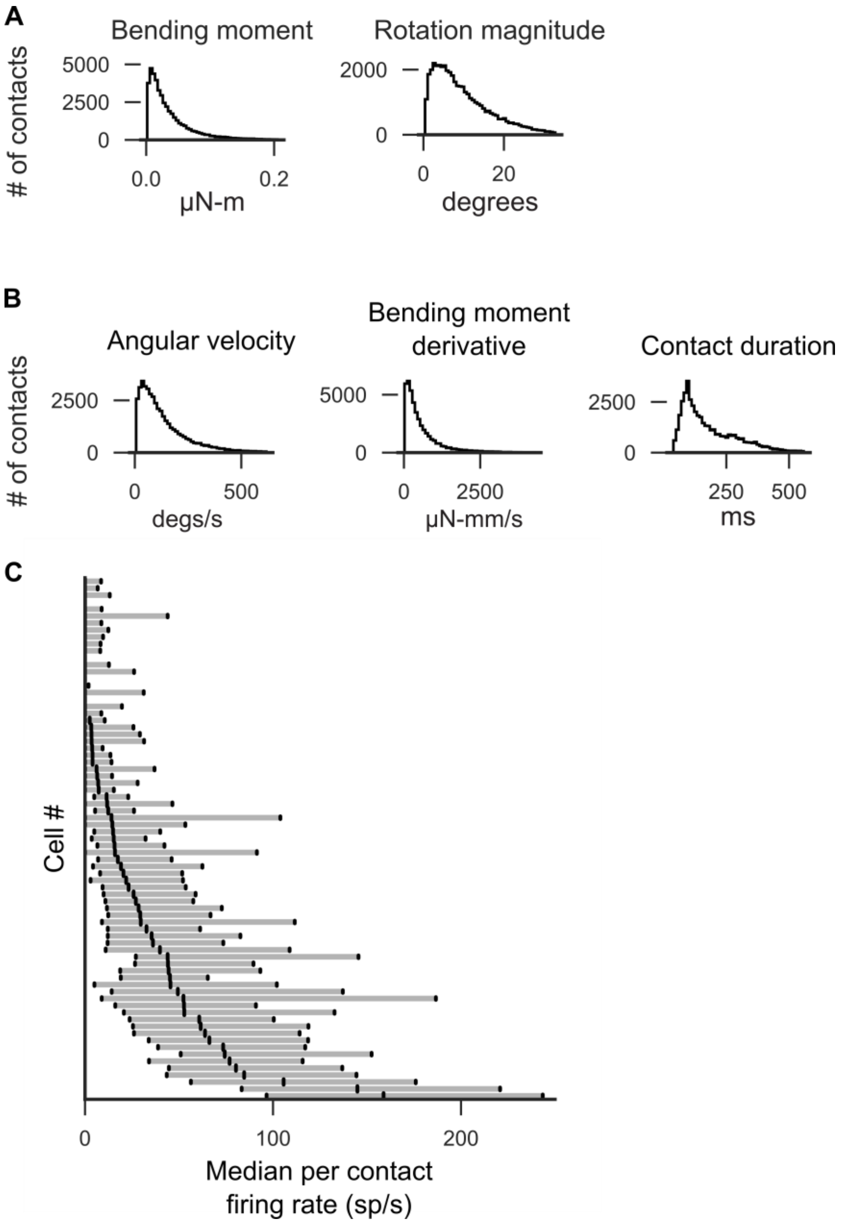


**Supplementary Figure 2: Marginal distributions of physical inputs and summary firing rates. (A)** Rotation magnitude and bending moment are computed as in *Methods.* Marginal distributions of these stimulus features are shown. The peak value reached during each contact is counted as the value for that contact. The histograms show the distribution of peak values across all contacts and all whiskers. **(B)** The trajectory of bending moment and rotation magnitude is followed for each contact until a peak value is reached. The peak value is divided by the time from onset of contact to peak value. These estimated derivatives are recorded as the angular velocity (rotation magnitude) and the bending moment derivative (bending moment) for that contact. The histograms of these derivatives over all contacts and all whiskers are shown. **(C)** The average firing rate for each contact is calculated as the number of spikes recorded during contact divided by contact duration. Median (middle point) and +/-IQR (shaded region) across all contacts are shown for each recorded cell. Average number of contacts per cell=648.

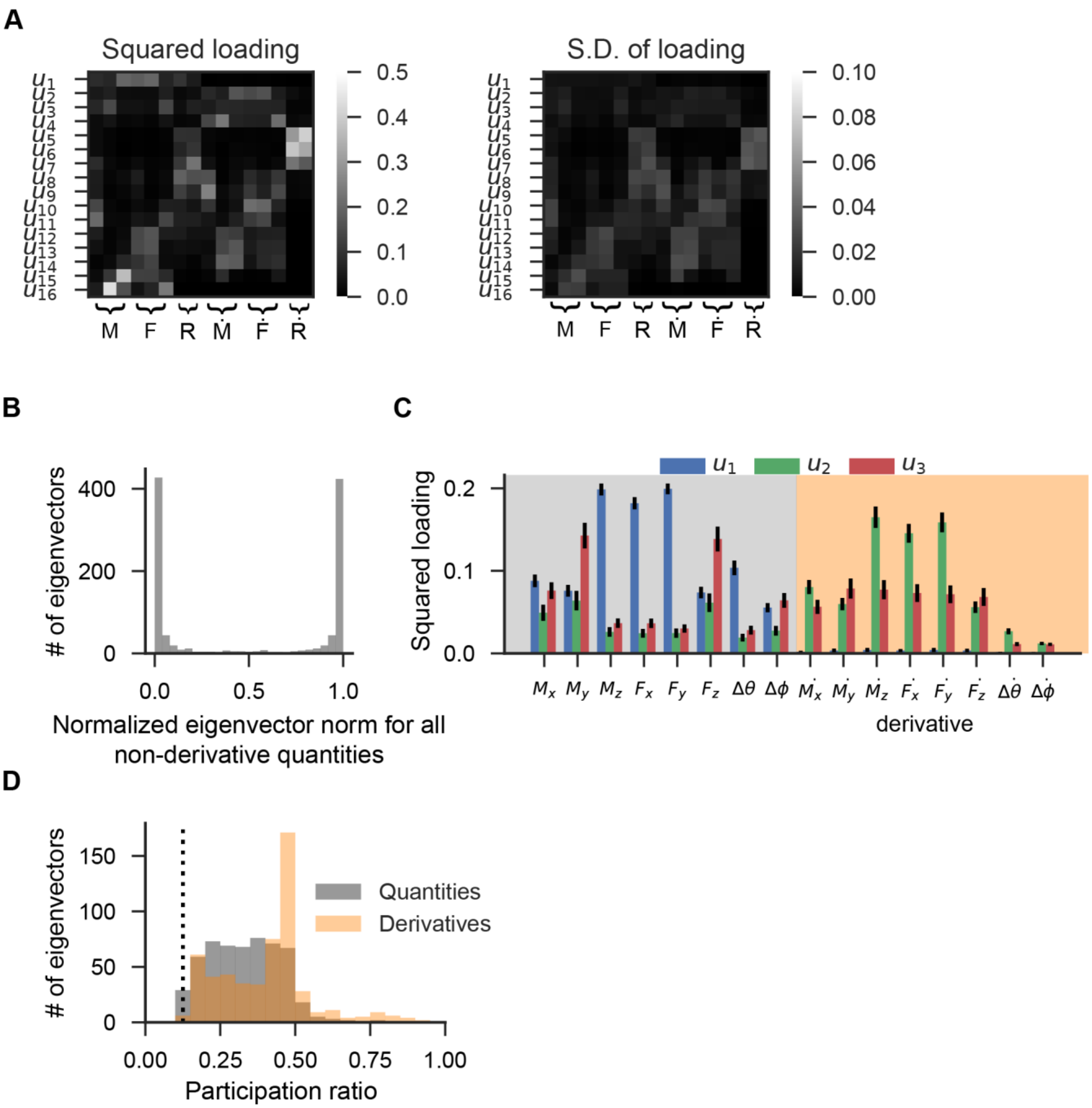


**Supplementary Figure 3: PCA descriptions of the input space.** PCA was applied to the observed mechanical features to obtain whisker specific low-dimensional representations of a 16-dimensional input space that incorporated components of moment, force, rotation, and their temporal derivatives (calculated as discrete difference after LOESS smoothing with a 95 ms window). **(A)** The squared loading of each mechanical component for the 16 PCs, shown as rows and ordered by variance accounted for. Shown left are averages across all reported whiskers; shown right are the standard deviations of the squared loadings across whiskers. **(B)** Histogram of the ratio of the L2 norm of the first half of each eigenvector, which comprises only the eight non-derivative quantities, to the L2 norm of the full eigenvector. The histogram includes all 16 eigenvectors for each measured whisker. The distribution peaks at 0 and 1, indicating that each eigenvector had significant loads on non-derivative or derivative quantities, but not on both. **(C)** Squared loading of each input component for each of the three leading eigenvectors, averaged across whiskers and labeled by the corresponding mechanical input variable. Error bars are +/- S.E.M. **(D)** The participation ratio is computed for each eigenvector, 16 per whisker, and reported for all whiskers. Eigenvectors were categorized as representing physical quantities (non-derivative) or their derivatives based on whether the summed norm of the squared loadings over all non-derivative quantities exceeded 0.5. Dashed line indicates a lower bound to the participation ratio, realized when all input components are equally weighted.

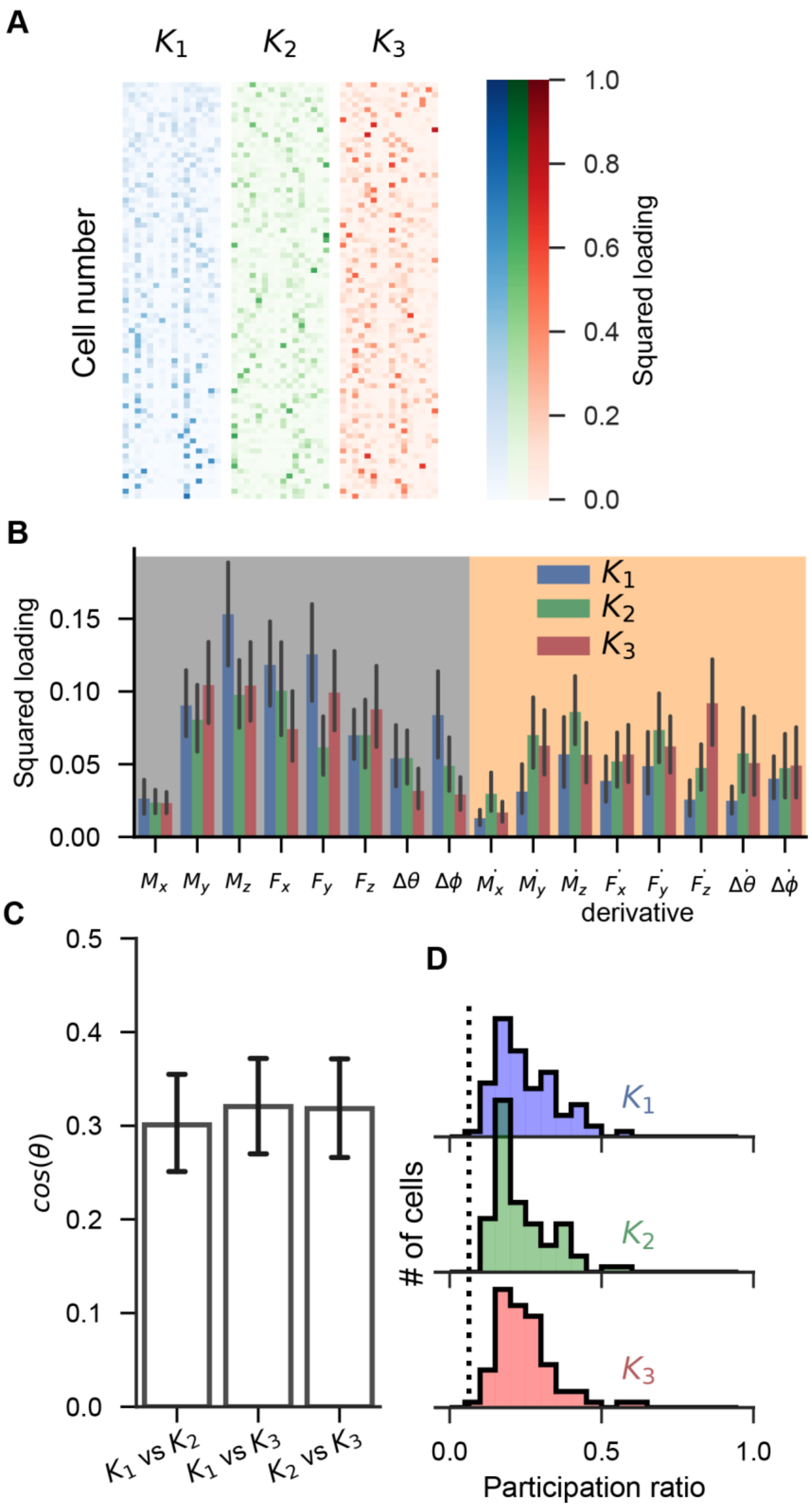


**Supplementary Figure 4: Neural representations of the input space.** Error bars are 95% confidence intervals in all panels. **(A)** Visualization of the GLM weights that characterize the response of all recorded Vg neurons. The response of each neuron is characterized by three vectors $\{K_i\}, 1 \leq i \leq 3,$ each a 16-dimensional vector with components associated with each of the input components. Each row is a neuron, sorted by the participation ratio for the first vector, $K_1$. Color represents each of the three vectors; color saturation represents the square of the corresponding component of the normalized $K$ vector. **(B)** Squared loadings of each input component for each of the normalized neural vectors, averaged across neurons and labeled by the corresponding mechanical input variable. **(C)** Overlap between normalized neural vectors; the pairwise cosines are shown, averaged across neurons. A value of zero indicates that the neural vectors are orthogonal; a value of 1 indicates collinearity. **(D)** Histograms of the participation ratio for each neural vector, for all neurons. Dashed line indicates a lower bound to the participation ratio, realized when all components are equally weighted. A participation ratio of 1 indicates that only one component of the neural vector is nonzero, indicating that only one of the input variables affects the firing rate of that neuron. In contrast, these results indicate that Vg neurons fire in response to distributed combinations of input variables.

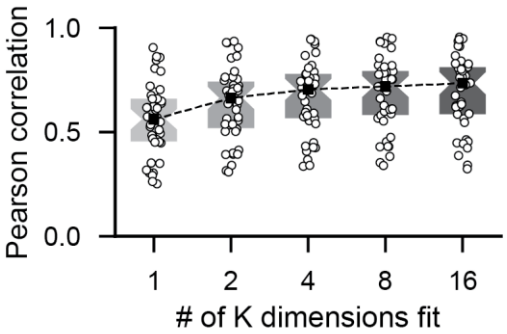


**Supplementary Figure 5: Performance of the neural model as a function of the number of neural vectors. (A)** Increasing the number of neural vectors increases model performance, which saturates at about four neural vectors in the full 16-dimensional input space. The Pearson correlation R is measured between the spike rate predicted by the model and the observed spike train smoothed with a gaussian kernel with $\sigma = 32$ ms. Boxes are median +/- 1 quartile.

**Supplementary Video 1:**

Whisker reconstruction, mechanics, and neural responses for example cell 1

*Top left*: Front-on view of whisker deflection. *Top right*: Top-down view of whisker deflection. *Bottom left*: 3D whisker reconstruction. Base point is shown in red and shifted to the origin. History of contact point from the last ten frames is shown along the whisker when contact occurs. *Bottom right*: Mechanical components (moments: orange; forces: cyan; rotations: purple) and spiking (vertical bars) elicited by corresponding whisker motion. Flicker of tip position is a result of failing tracking and is ignored in all analyses.

**Supplementary Video 2:**

Whisker reconstruction, mechanics, and neural responses for example cell 2

*Top left*: Front-on view of whisker deflection. *Top right*: Top-down view of whisker deflection. *Bottom left*: 3D whisker reconstruction. Base point is shown in red and shifted to the origin. History of contact point from the last ten frames is shown along the whisker when contact occurs. *Bottom right*: Mechanical components (moments: orange; forces: cyan; rotations: purple) and spiking (vertical bars) elicited by corresponding whisker motion. Flicker of tip position is a result of failing tracking and is ignored in all analyses.

**Supplementary Video 3:**

Whisker reconstruction, mechanics, and neural responses for example cell 3

*Top left*: Front-on view of whisker deflection. *Top right*: Top-down view of whisker deflection. *Bottom left*: 3D whisker reconstruction. Base point is shown in red and shifted to the origin. History of contact point from the last ten frames is shown along the whisker when contact occurs. *Bottom right*: Mechanical components (moments: orange; forces: cyan; rotations: purple) and spiking (vertical bars) elicited by corresponding whisker motion. Flicker of tip position is a result of failing tracking and is ignored in all analyses.
